# RNA activities of trans species: RNA from the sperm of the father of an autistic child programs glial cells and behavioral disorders in mice

**DOI:** 10.1101/2023.10.27.564312

**Authors:** Zeynep Yilmaz Sukranli, Keziban Korkmaz Bayram, Ecmel Mehmetbeyoglu, Zuleyha Doganyigit, Feyzullah Beyaz, Elif Funda Sener, Serpil Taheri, Yusuf Ozkul, Minoo Rassoulzadegan

## Abstract

Recently, we described the alteration of six miRNAs in the serum of autistic children, their fathers, mothers, siblings and in the sperm of autistic mouse models. Studies in model organisms suggest that noncoding RNAs participate in transcriptional modulation pathways. Using mice, approaches to alter the amount of RNA in fertilized eggs enable *invivo* intervention at an early stage of development. Noncoding RNAs are very numerous in spermatozoa. Our study aims to address a fundamental question: Can the transfer of RNA content from sperm to eggs result in changes in phenotypic traits, such as autism? To explore this, we utilized RNA from the sperm of a father with autistic children and microinjected it into fertilized mouse eggs, thus creating mouse models for autism. Here, we induced in a single step by microinjection of sperm RNA into fertilized eggs the transformation of glial cells into cells capable of developing ‵autism-like‵ disorders in mice born from these alterations.

## Introduction

Autism is one of the most common behavioral changes detected early in life^1^. Despite extensive research, the etiology of autism remains very complex. The cause of autism is linked to the genetic and environmental history of parents which together could lead to the development of autism spectrum disorders (ASD) in children^2,3^. A recent survey reveals several hundred distinct sites of nucleotide changes in the genome between patients^4^. Several environmental and genetic factors identified in humans have already been validated with mouse models^5–6^, however, have not yet been associated in all autistic patients^7–8^. Evidence that the altered level of six miRNAs (miR-19a-3p, miR-361-5p, miR-3613-3p, miR-150-5p, miR-126-3p, miR-499a-5p) in parents and thus inheritance through the expression of altered phenotypes^9^ would lead to the inclusion of a common genetic marker for autism cases. Altered levels of RNAs such as non-coding RNAs can be passed on to the next generation via the germline to alter offspring phenotypes ^10,11^.

However, how RNA subornation is established and maintained throughout generation remains a matter of speculation. Recently, on the activity of sperm RNA characterized in mouse models, a growing number of experiments highlight it as an important source of parental hereditary information above DNA^12–14^. The six miRNAs listed above are present in sperm RNA in general and are altered in the father of an autistic child. Five of six (miR-19a-3p, miR-150-5p, miR-126-3p, miR-361-5p, miR-499a-5p) are also present in mouse sperm, the sixth miR-3613-3p does not exist in the mouse genome. These six inherited miRNAs are then found downregulated in the sera of autistic patients and ‵autism-like‵ mouse models. One of the questions posed here is to examine sperm RNA activities of that cover a possible paternally acquired autistic phenotype. Microinjection of sperm RNA into fertilized mouse eggs could lead to variable phenotypes depending on the origin of the males from which the RNA is collected^10^. In particular, although exposure of total RNA from (human) sperm to fertilized mouse eggs may have adverse effects on offspring, can symptoms of autism also become established in the next generation? Here, we focus on experimentally induced autistic traits, particularly the potential of human sperm RNAs to induce “autism-like” phenotypes in mice.

Offspring born from eggs micro-injected with sperm RNAs from the father of the autistic child’s were compared to controls at the molecular level and by behavioural assessment (open field maze, recognition of a familiar and novel object, social interaction, tail suspension, Marble burying and elevated plus maze assays).

In addition, we examined the prefrontal cortex, corpus callosum, striatum, hippocampus, amygdala and cerebellum, which have been implicated in the pathophysiology of autism in the brain^15,16^. Our study involved, the examination of the brain anatomy in animals from all groups, utilising immuno-histochemical stains. Notably, differences were observed in animals induced by valproic acid (VPA) and microinjected RNA. While astrocytes showed abnormal astrogliosis (and microgliosis, which is normally invaded by microglia), no signs of astrocyte disorientation or microglial extensions which are often seen in the autistic human brain, were detected. Moreover, we did not observe hypertrophy GFAP and Iba-1 positive-cells or hyperplasia were detected in immunopositive cells.

Microglia and astrocytes play important roles in the regulation of neuronal metabolism, providing nutritional support of neurons, participating in the synapses formation and facilitating neurotransmission^4,6,17,18^. Furthermore, in the microinjected with RNA group, there was a significant increase in the number of oligodendrocyte products accompanied by branching, compared to the controls groups. These results could be attributed to the observed autistic traits in both groups.

In appears that in both groups of mice the manifestation of autistic behavioral disorders may be linked glial cell disorders.

## Results

### RNA targeted autism-like mouse model

miRNAs have been found down-regulated in the blood of autistic children and in the mouse model of autism^9^. Among the Turkish cohort of 37 families including one or more children with behavior disorders (45 subjects altogether)^9^, we were interested in a family with two affected children. See Figure 2. A father from two separate marriages has multiple children and one of each child from these separate unions develops a distinct psychological disorder. We have previously shown that all of them (six miRNAs) are therefore already altered in their father’s sperm. We examined whether the RNAs of this human spermatozoa contain indicator RNAs? If so, could it also induce behavioral changes in adult mice after microinjection into fertilized mouse eggs? Humans and mice despite significant differences in DNA and gene expression, use a similar gene regulatory mechanism and network. DNA mutation linked to disease in humans often has homologs in the mouse genome. The major epigenetic changes known to date in rodents are found in humans. However, common epigenetic functions remain relatively understudied. To investigate the potential for transgenerational control of ASD in mice via RNA, we used the same approach previously used with mouse sperm RNA to evaluate with human sperm RNA.

In our functional studies in mice, we assume that exposure (after microinjection) of the genome of a fertilized mouse egg to sperm RNA from a mouse model of autism or from the father of an autistic child will lead to modifications in the expressions of autism-related miRNAs transcripts in blood and adult mouse tissues. To begin, we selected sperm RNA from a valproic acid treated (VPA) autistic mouse model (previously characterized in our laboratory) for functional analysis in mice see Table-S2. We also chose RNA from the sperm of a father of an autistic child because the six miRNAs, as reported in our prior study^9^ exhibited a significant downregulation (50% of controls) in the parents’ blood of the patients with autism. Furthermore, these miRNAs were found to be altered in an available sperm sample from one of the fathers of two affected children.

Variations in transcripts play an essential role in brain development^19,21,22,24^. Therefore, when establishing an RNA–induced autism-like mouse model we exposed fertilized mouse eggs to exogenous total RNA from sperm (microinjected RNA see Figure 1 for experimental plan). As expected, F0 (F0-spRNAs) males exhibited transcriptional and phenotypic variations compared to control mice.

**Figure 1.**
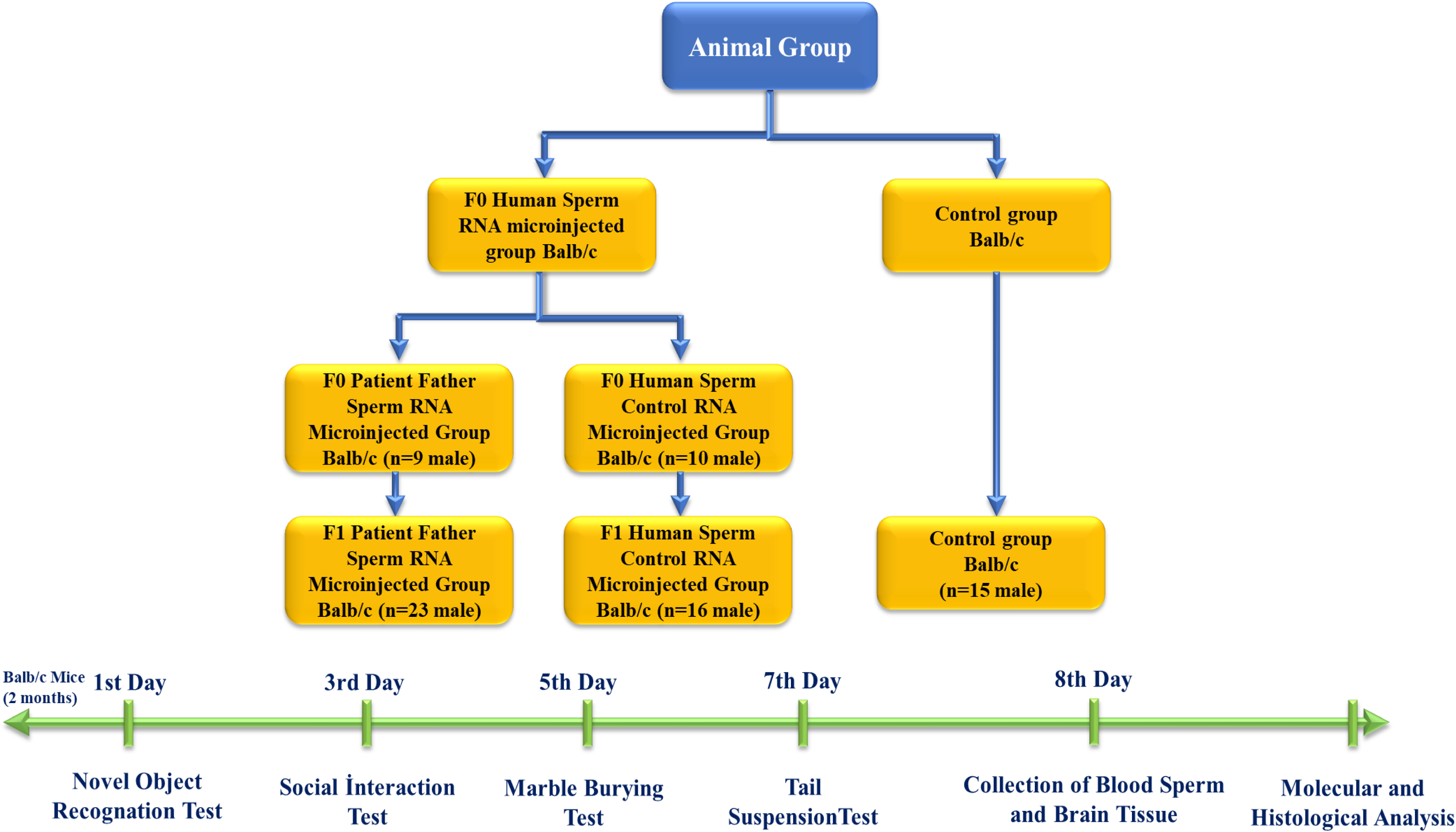
Experimental design and timeline as graphical abstract

**Figure 2.**
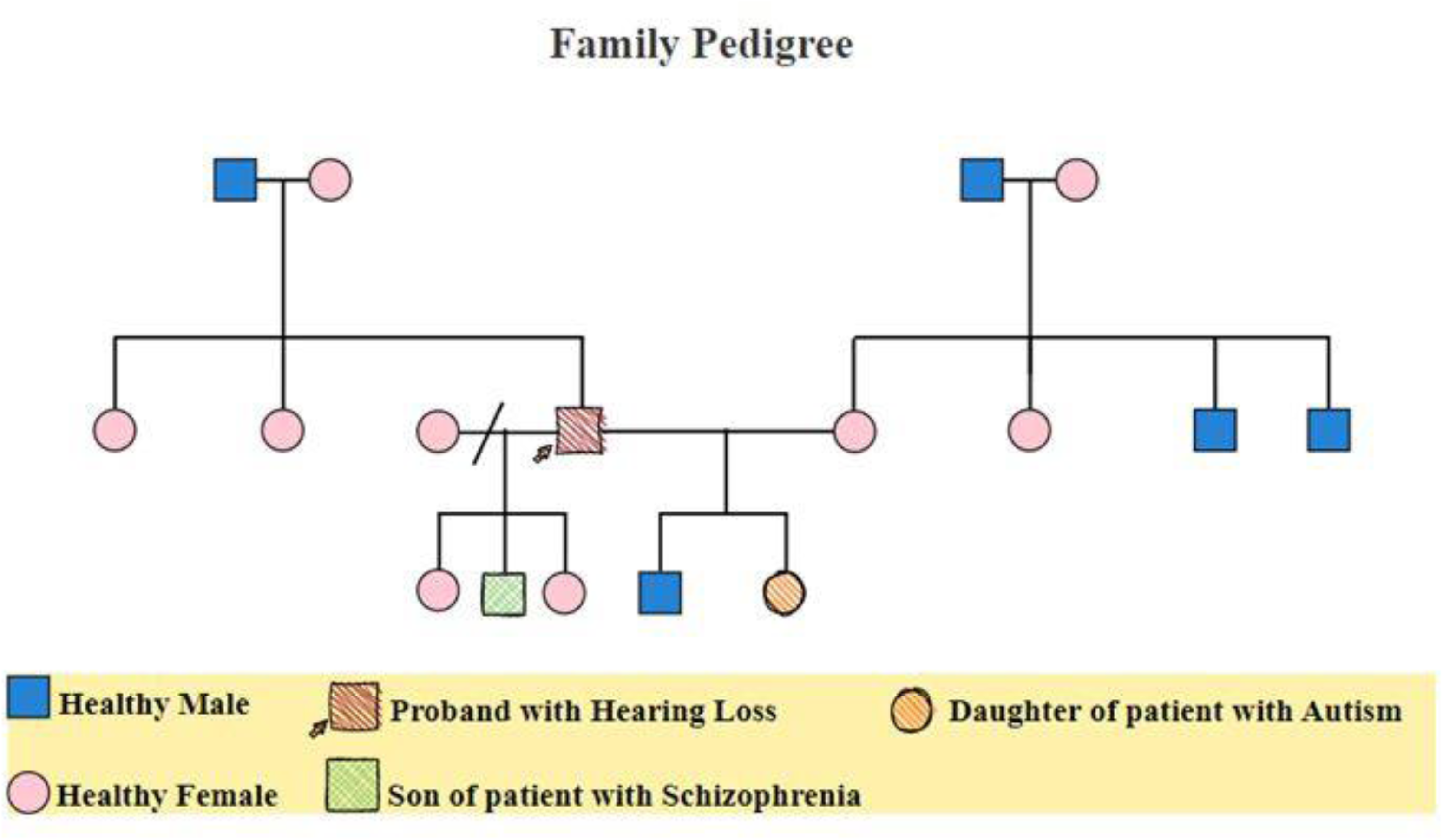
Illustration of family pedigree of patient whose father sperm RNA was microinjected into mouse embryos

Previous analysis of six miRNAs (−5p and 3p strands) from the father’s sperm showed that all were modified. Mice born after microinjection of paternal sperm RNA again showed significant downregulation of four miRNAs 361-3p, 150-5p, 126-5p, and 499-3p. Paternal sperm RNA downregulates at least four miRNAs in the blood of adult mice (Figure 4). The two miRNAs up-regulated in paternal sperm 361-3p (reverse of 5p) and 499-3p (same forward) are now downregulated in mice. Two miRNAs 361-5p and 499-3p upregulated in father’s sperm (see Figure 5, Supplemantary Table 2). This may suggest that upregulation in sperm triggers downregulation in the next generation.

#### Behavioral variations

Genes linked to ASD are also found in mice and have been analyzed by targeted mutagenesis. Basically, so far, all genetic mutations linked to ASD also perturbs brain development when tested in mice (Review Crawlay^20^). Although all the deficits defining autism are difficult to assess in mice, significant efforts are being made to develop behavioural tests to phenotype mice. Based on the behavioral neuroscience literature we examined mouse colonies on existing behavioral tasks to refine key symptoms. The summary schedule for the behavioral task analysis is listed in Figure 1.

Behavior tasks were carried out on male mice from the age of 2 months by recognition of a novel versus familial object, social interactions, suspension by the tail, marble burying assays (see timeline). While it was observed that the interest in the new object was higher in all groups in the novel object recognition test, no significant difference was detected between the groups in terms of discrimination index, total movement and speed. In the social interaction test, all groups, except the control group, including the F0 and F1 generations, showed more interest in the empty cage and exhibited behaviours that did not support social life and lacked empathy. In the Marble test, there is an increased marble burying activity towards the next generation compared to the control, which is an indicator of hippocampal damage. This situation is clearly observed in the patient sperm RNA group. In the tail suspension test, where anxiety findings were measured, it was determined that although the most inactive group was the F1 human sperm RNA control group, all groups exhibited similar phenotype behaviour to the control.

### Histological examinations

The prefrontal cortex, corpus callosum, striatum, hippocampus, amygdala and cerebellum, are widely recognized as regions associated with autism. In all groups brains were studied in detail with histochemical stains such as hematoxylin-Eosin and Luksol fast Blue. The structure of the corpus callosum in the brains of all groups of animals was examined in the striatum and hippocampus regions and found to have a normal structure, as in the brains of animals in the control group. No agenesis of the corpus callosum was reported in any of the experimental groups observed in the brains of autistic individuals (Figures 6 and 7). The lateral ventricles and third ventricle were examined in the brains of all groups of animals and the dimensions of the ventricles were determined to be normal as in the brains of the control group animals. In other words, ventricular dilation, which is believed to be present in the autistic human brain, was not observed in animals from all experimental groups.

At both anatomical and histological levels, no specific changes were observed in the cerebellum and brainstem when comparing animals in all experimental groups to those in the control group. Furthermore, there was an absence of white matter substances between the hemispheres in the region of the striatum and the hippocampus along the midline in all groups.

It was observed that the outer capsule, inner capsule, and anterior commissure in the brains of all groups of animals did not present a different status from that of the brains of animals in the control group. It was determined that the sizes of the striatum and septum as well as the cortical thicknesses of animals in all groups did not differ from those of control mice.

No differences were observed in the brains of all groups of mice compared to the brains of control animals in layers CA1, CA2, CA3 and from the dentate gyrus to the hippocampus. Similarly, no discernible distinctions were noted in the anatomical and cellular structure of amygdalin in the brains of all groups of mice compared to the brains of animals in the control group. Furthermore, there was no reduction in crescent size of the hippocampal in the brains of all groups of mice.

At the cellular level, neurodegeneration was not observed in the prefrontal cortex, striatum, hippocampus and cerebellum regions of animal brains in all groups, however gliosis was observed in the VPA, Human sperm-RNA and reverse miRNAs groups. No ectopic white matter clusters, as reported in autistic human brains in singular cortexes and other cortexes, were found in the brains of all groups of mice. Histological examination of the cerebellum in all groups of mice did not reveal areas of Purkinje cell loss which has been reported in the autistic human cerebellum (Figure 3).

**Figure 3.**
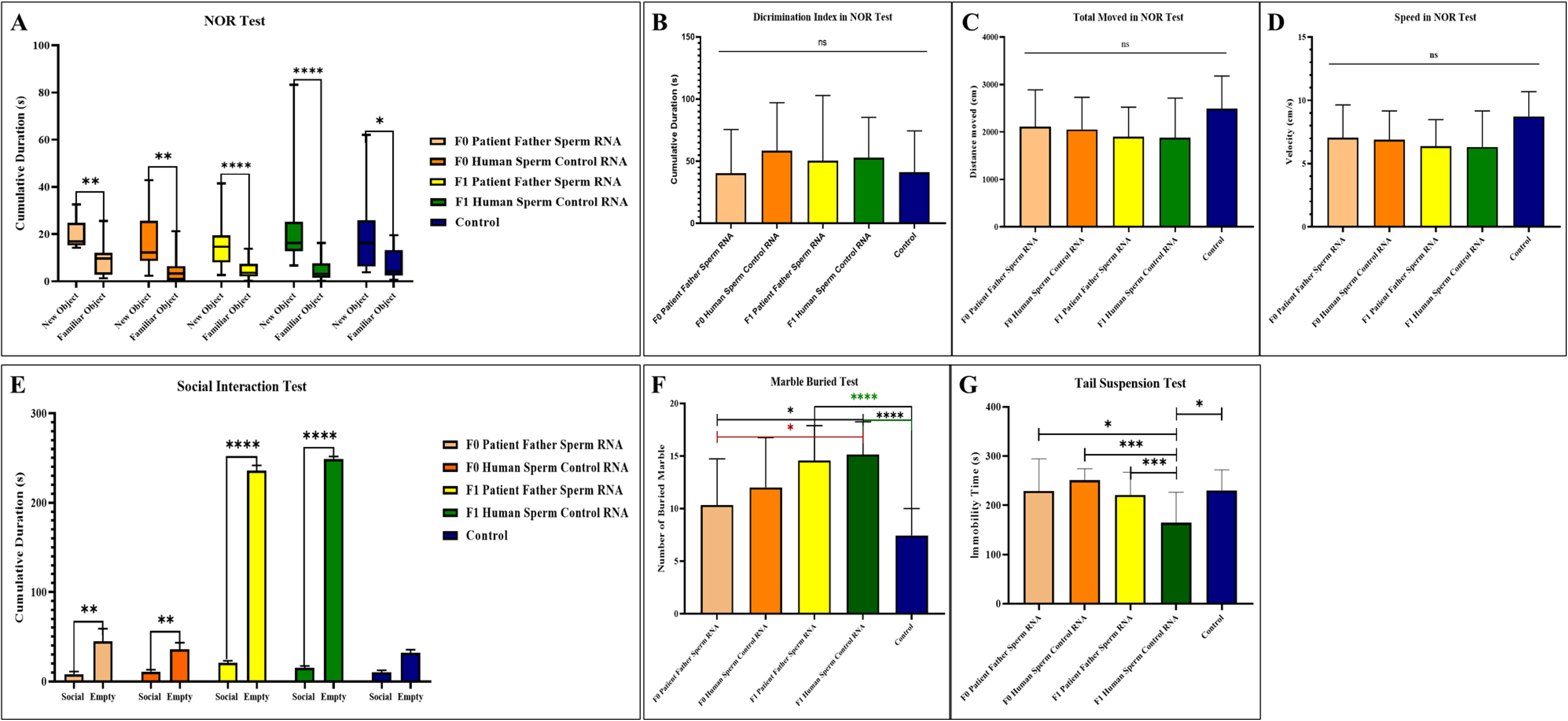
Behavioral experiment results performed in adulthood of mice to which patient father sperm RNA and healthy sperm RNA were transferred by microinjection. **A.** Measuring the interest of mice to the new object in the novel object recognition test, **B.** Novel object recognition (NOR) test showing the difference in terms of the discrimination index between groups, **C.** Demonstration of the distance traveled by mice during the novel object recognition test, **D.** Demonstration of the speed of mice during the novel object recognition test, **E.** Demonstration of mice’s social interest and interest in the empty cage during the social interaction test, **F.** Display of the number of marbles buried by mice in the Marble Buried Test, **G.** Illustration of the time spent inactive by mice in the tail suspension test.

**Figure 4.**
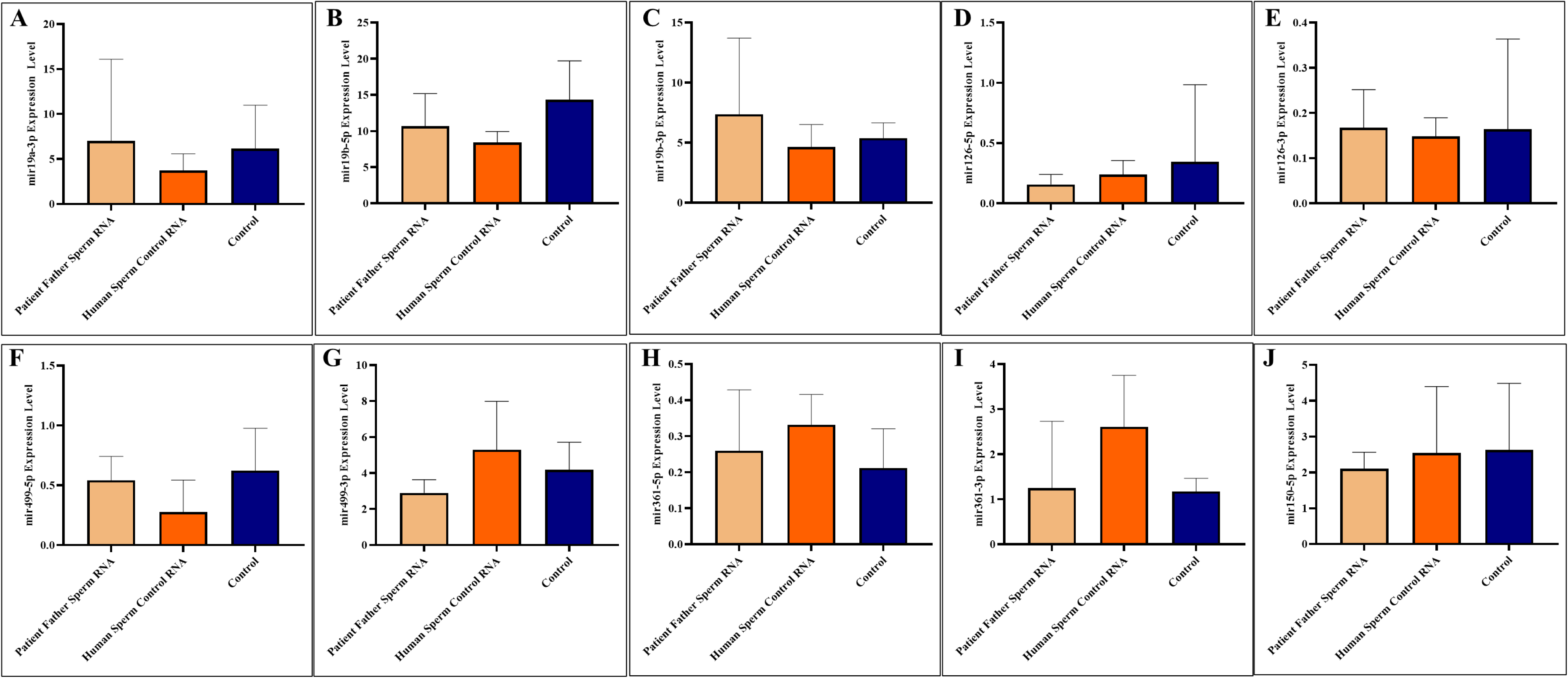
Expression levels analysis of miR-19a-3p, miR-19b-5p, miR-19b-3p, miR-126-5p, miR-126-3p, miR-499a-5p, miR-499a-3p, miR-361-5p, miR-361-3p, and miR-150-5p in the blood of mice that received patient father sperm RNA and healthy sperm RNA through microinjection. Blood samples were collected upon sacrifice in adulthood (A-J).

### Immunohistochemistry

The prefrontal cortex, corpus callosum, striatum, hippocampus, amygdala and cerebellum, suspected to be associated with the pathophysiology of autism in the brains of animals of all groups, were studied by immuno-staining-histochemical. Differences were observed in the VPA, miR-reverse and total-RNA groups. Astrocytes displayed abnormal features characterised by astrogliosis (and microgliosis, which is normally invaded by microglia). However, it is important to note that no instances of astrocyte misorientation or microglial extension as reported in the human brain with autism, were observed. Furthermore, there was no evidence of hypertrophy or hyperplasia of GFAP and Iba-1 cells in immunopositive cells (Figure 11-13).

Microglia cells and astrocyte are known to play an important role in regulating neuronal metabolism, providing nutritional support to neurons, facilitating synapse formation and contributing to neurotransmission. Furthermore, in the total sperm human RNA group (from both the father of autistic children and the normal group), the number of oligodendrocyte products was increased and significantly branched compared to the control and VPA groups. So, at this point, these results reveal an additional side effect of human sperm RNAs rather than just autism. Unlike the miRs previously examined, 3-p miR-RNAs also induce the same changes as observed in sperm total RNAs and VPA groups in astrocytes and microglia. Although miR-19a and miR-499b 5-p did not produce the same effects. When examining parental sperm RNAs, all three groups are exhibited behavioural disorders such as autism-like disorders comparable to the VPA groups.

## Discussion

The present study revealed the regulatory function of sperm RNA in different species including that of sperm RNA derived from the father of an autistic child. Through a program capable of modifying the phenotypes of glial cell and interfering with the mouse development, it was possible to establish an ‵ASD-like‵ mouse model. The analysis of autism-related miRNAs transcripts revealed a down-regulation in affected glial cells accompanied by changes in behavioural phenotypes (Figure. 5). These results suggest that RNAs variations may contribute to the development of ASDs. In maize, multiples cases of small RNAs variations responsible for changing alleles- in trans with phenotypic changes in the next generation have already been reported ^23^. Furthermore, there was a correlation between the effects induced by valproic acid (VPA) and those induced by human sperm RNA, suggesting shared pathways affected by both the mouse VPA model and sperm RNA (human) induced phenotype. Although glial cells are thought to play a major role in brain development and pathogenesis in general, concrete evidence for their initiating role in autism is still lacking, as such analyses are typically conducted post mortem. It is important to pay attention to developmental patterns early on as they may offer valuable insights into ASD symptoms. In particular, our study in mice suggest that glial cells are the first affected. A recent review clearly highlights different functions of glial cells in autism^8^.

**Figure 5.**
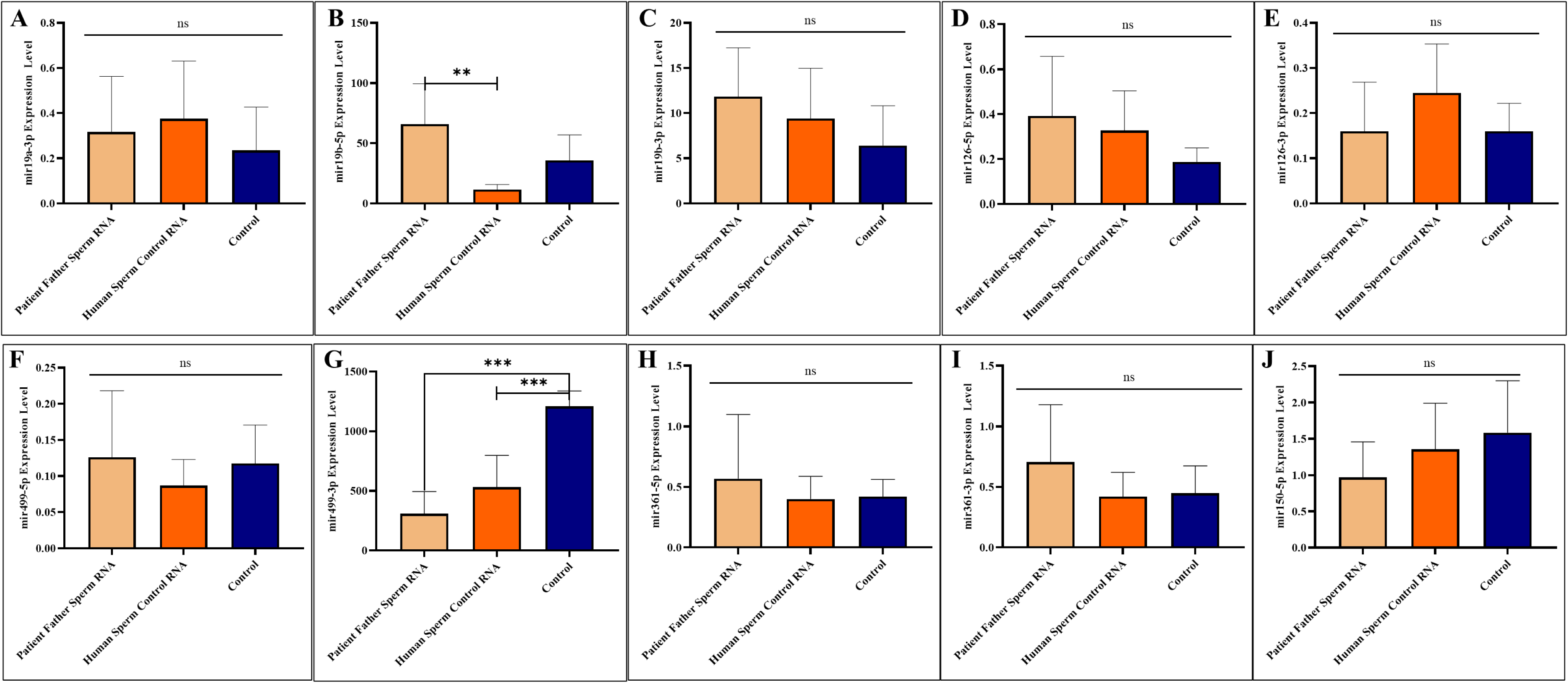
Expression levels analysis of miR-19a-3p, miR-19b-5p, miR-19b-3p, miR-126-5p, miR-126-3p, miR-499a-5p, miR-499a-3p, miR-361-5p, miR-361-3p, and miR-150-5p in the sperm of mice that received patient father sperm RNA and healthy sperm RNA through microinjection Sperm samples were collected upon sacrifice in adulthood (A-J).

**Figure 6.**
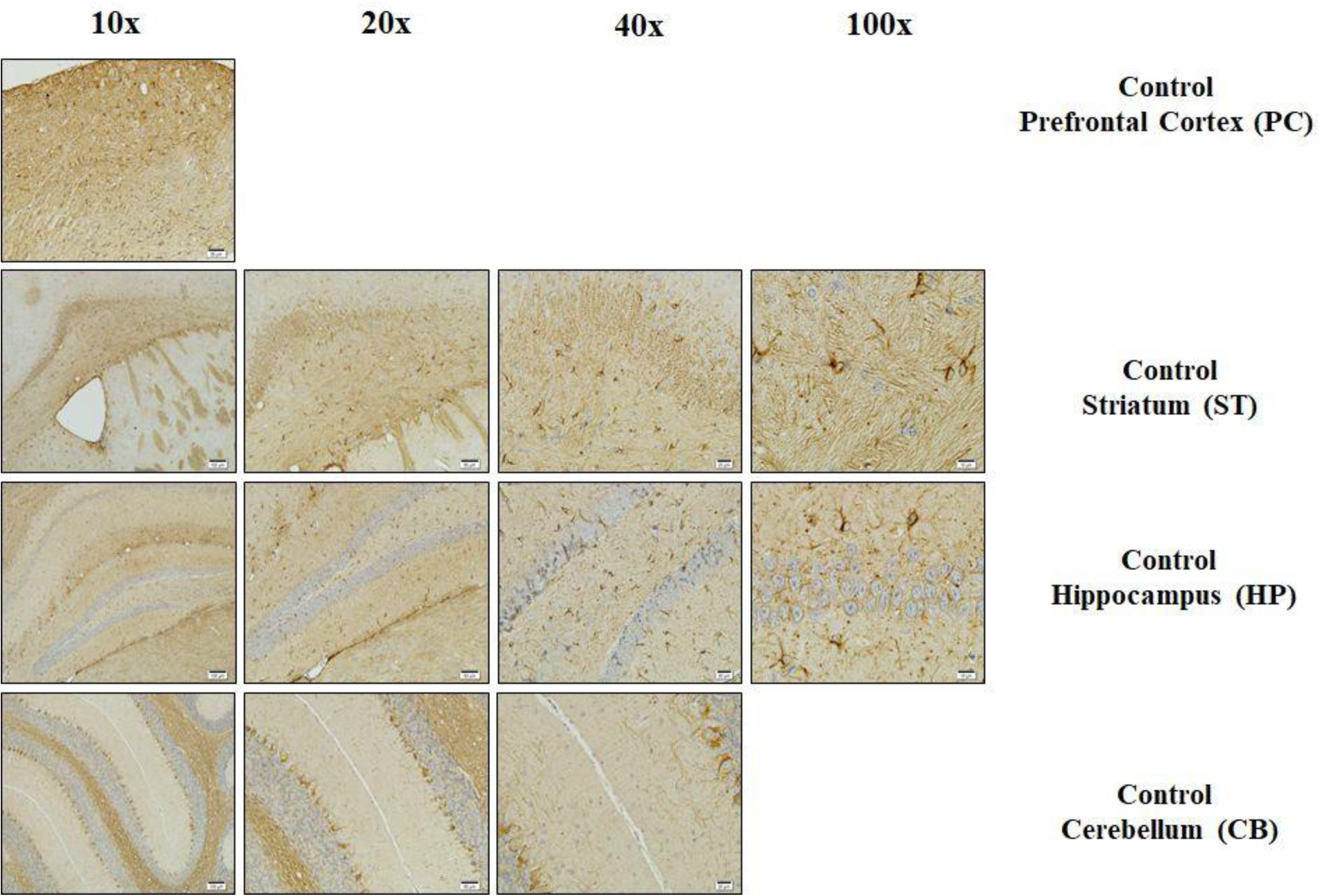
The control group. Histological examination of the prefrontal cortex, striatum, hippocampus and cerebellum brain regions:

**Figure 7.**
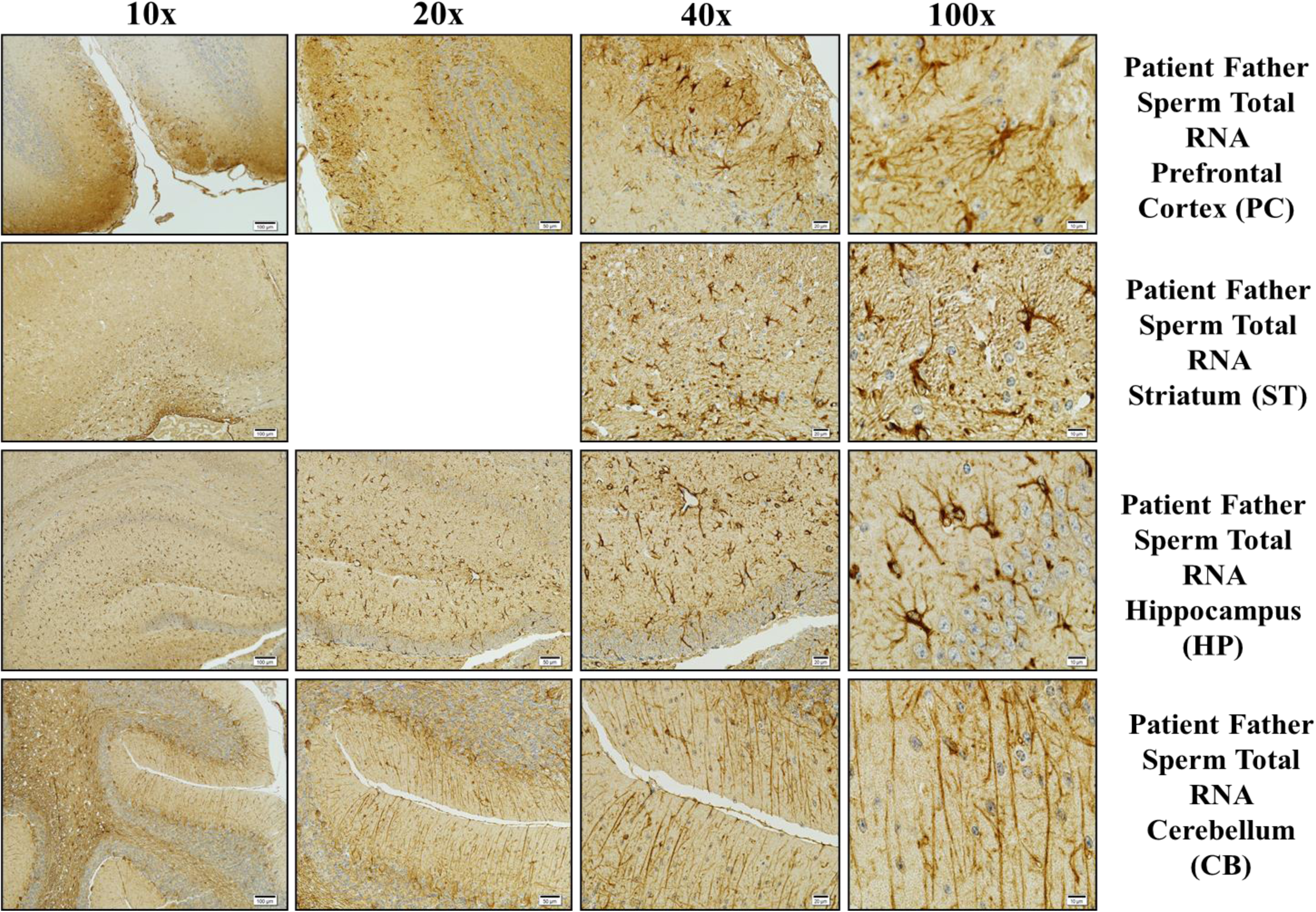
The patient father sperm RNA group

**Figure 8.**
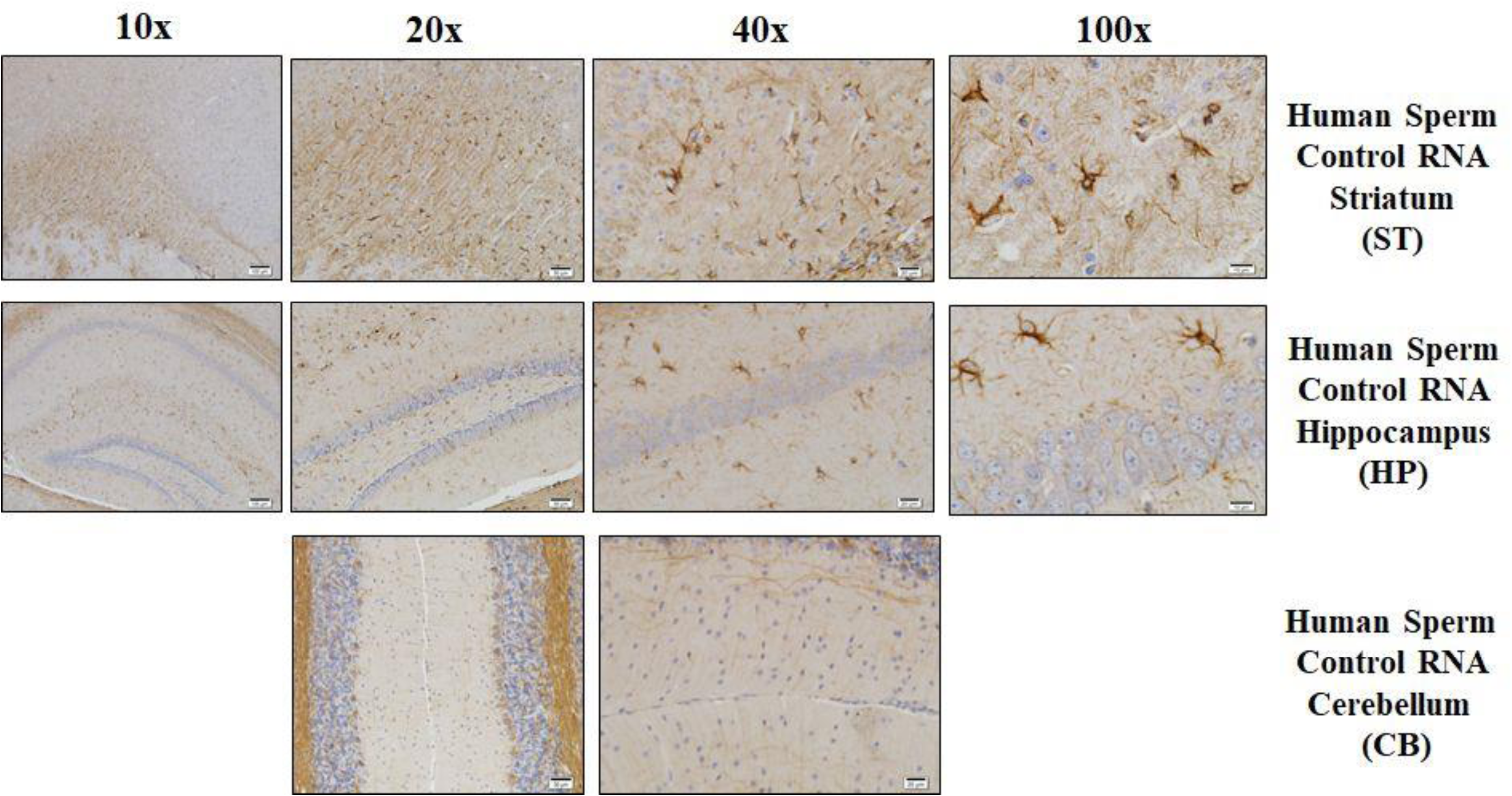
The human sperm control RNA group. Fluorescent staining in the hippocampus, cerebellum and striatum brain regions:

**Figure 9.**
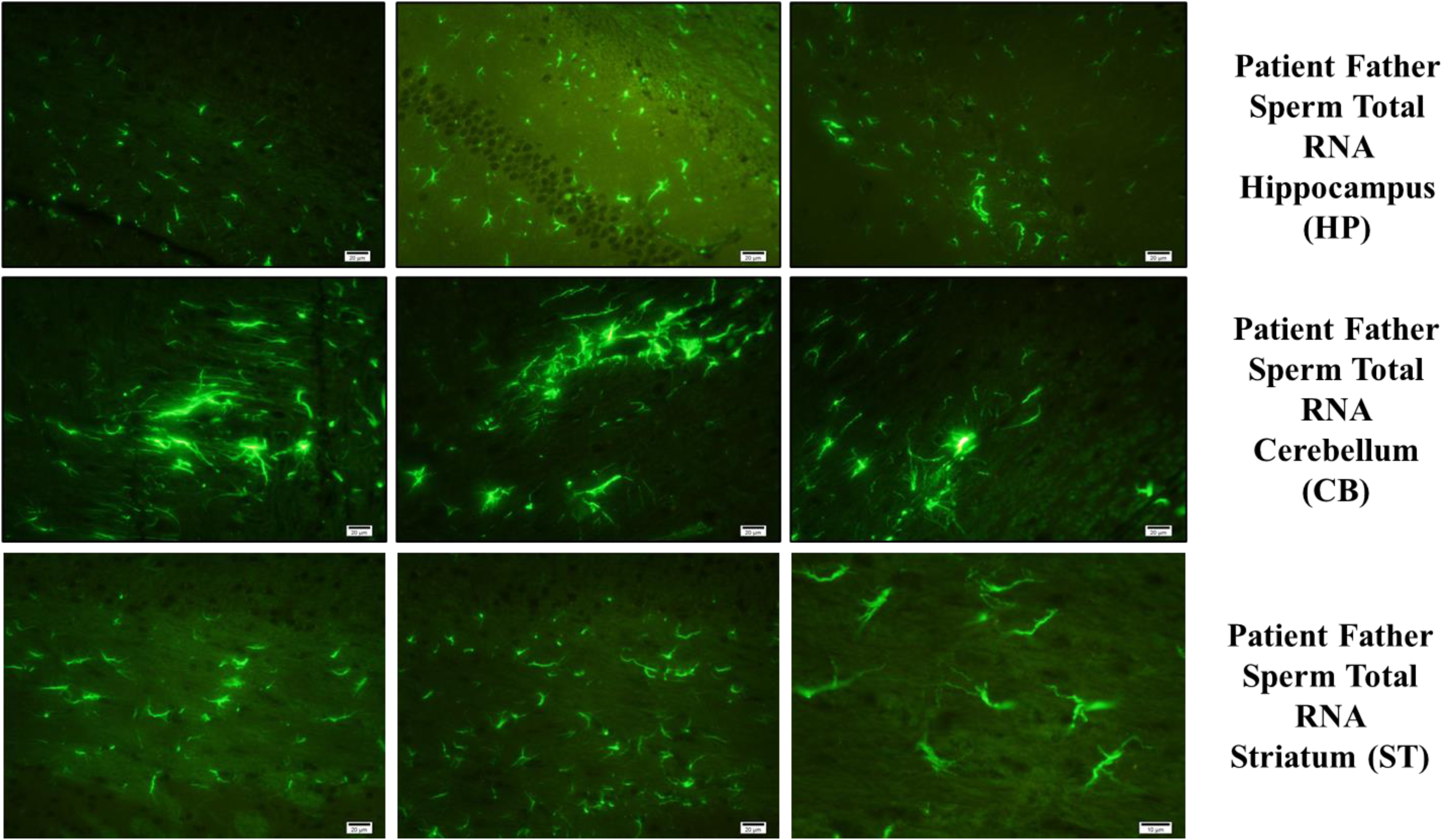
The patient father sperm total RNA microinjected group.

**Figure 10.**
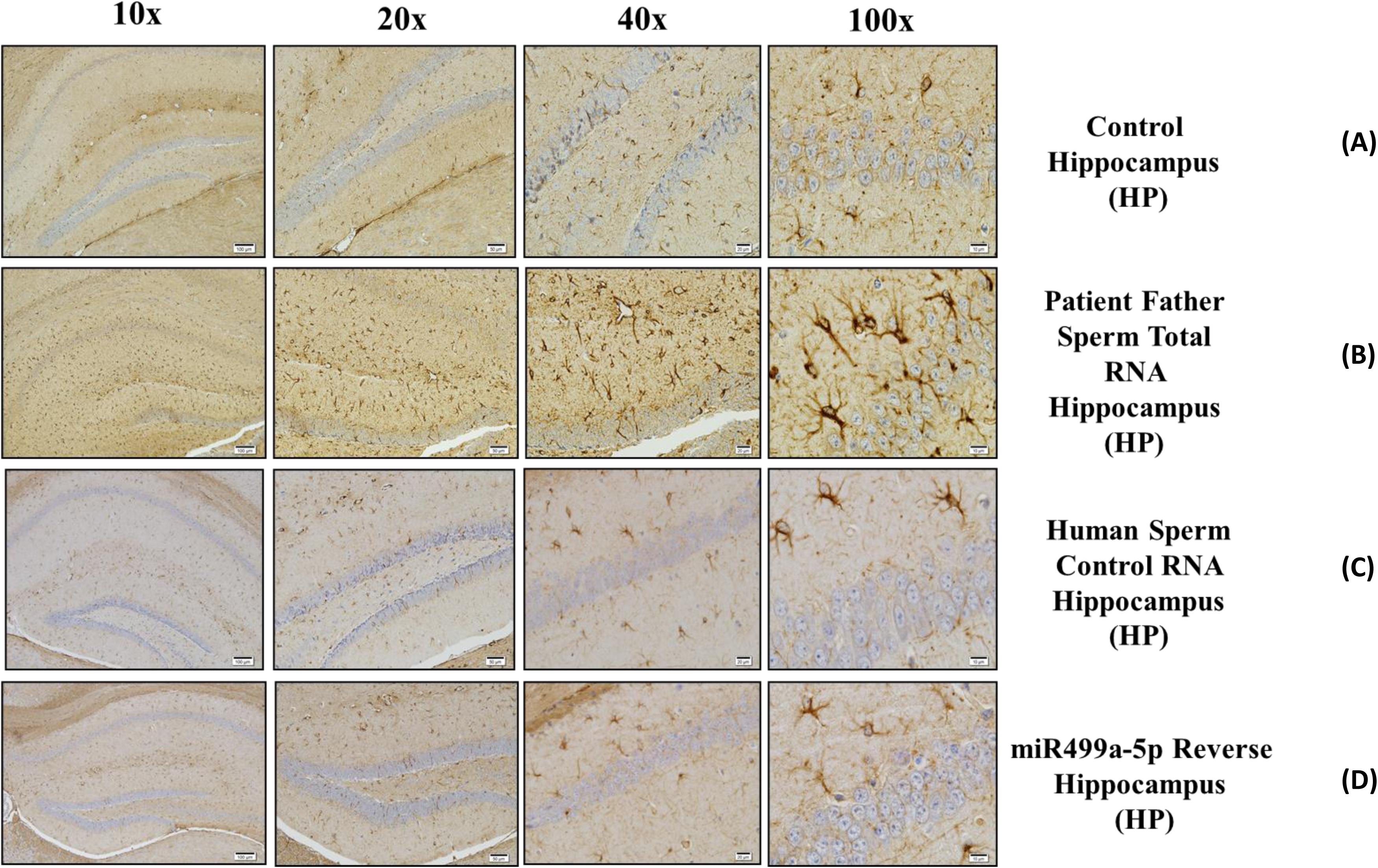
Histological examination of the hippocampus in different groups **A.** Control group, **B.** Patient father sperm RNA group, **C.** Human Sperm Control RNA group and **D.** mir499a-5p reverse group.

**Figure 11.**
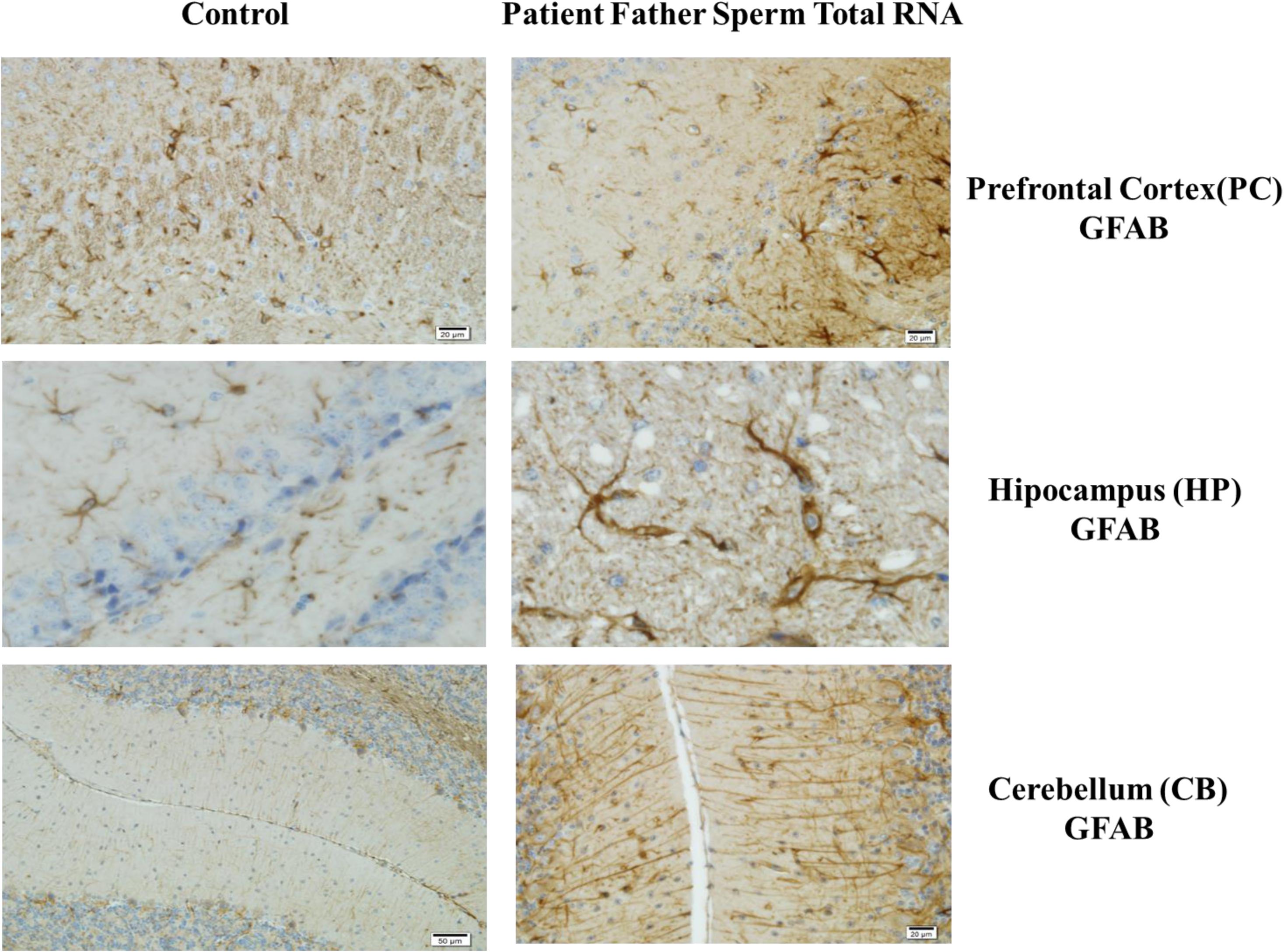
Visualization of expression levels in the prefrontal cortex, hippocampus and cerebellum with GFAB Ab staining and comparison of the patient father sperm RNA group with the control.

**Figure 12.**
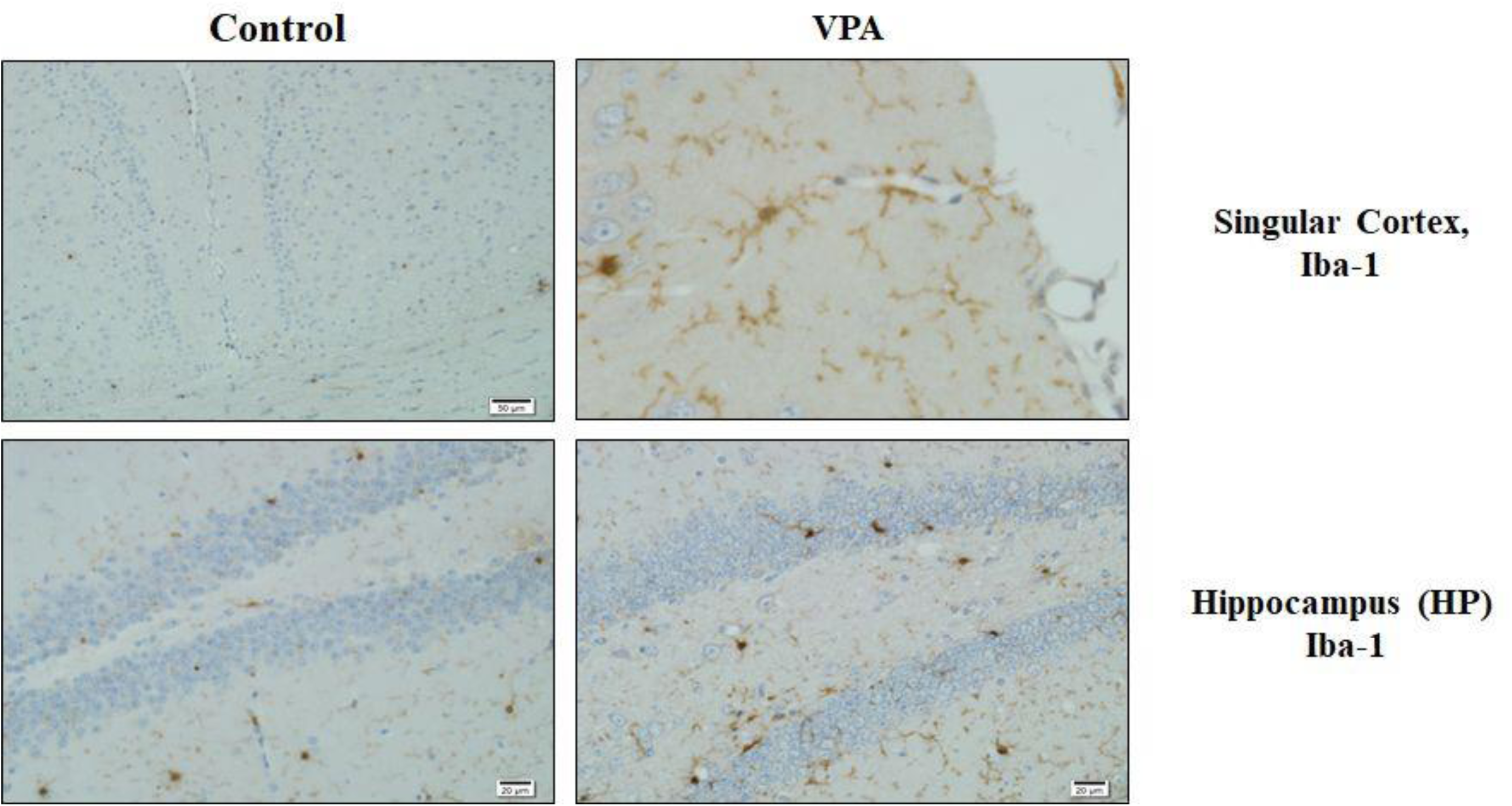
Visualization of expression levels in the singular cortex and with Iba-1 Ab staining and comparison of the VPA group with the control.

**Figure 13.**
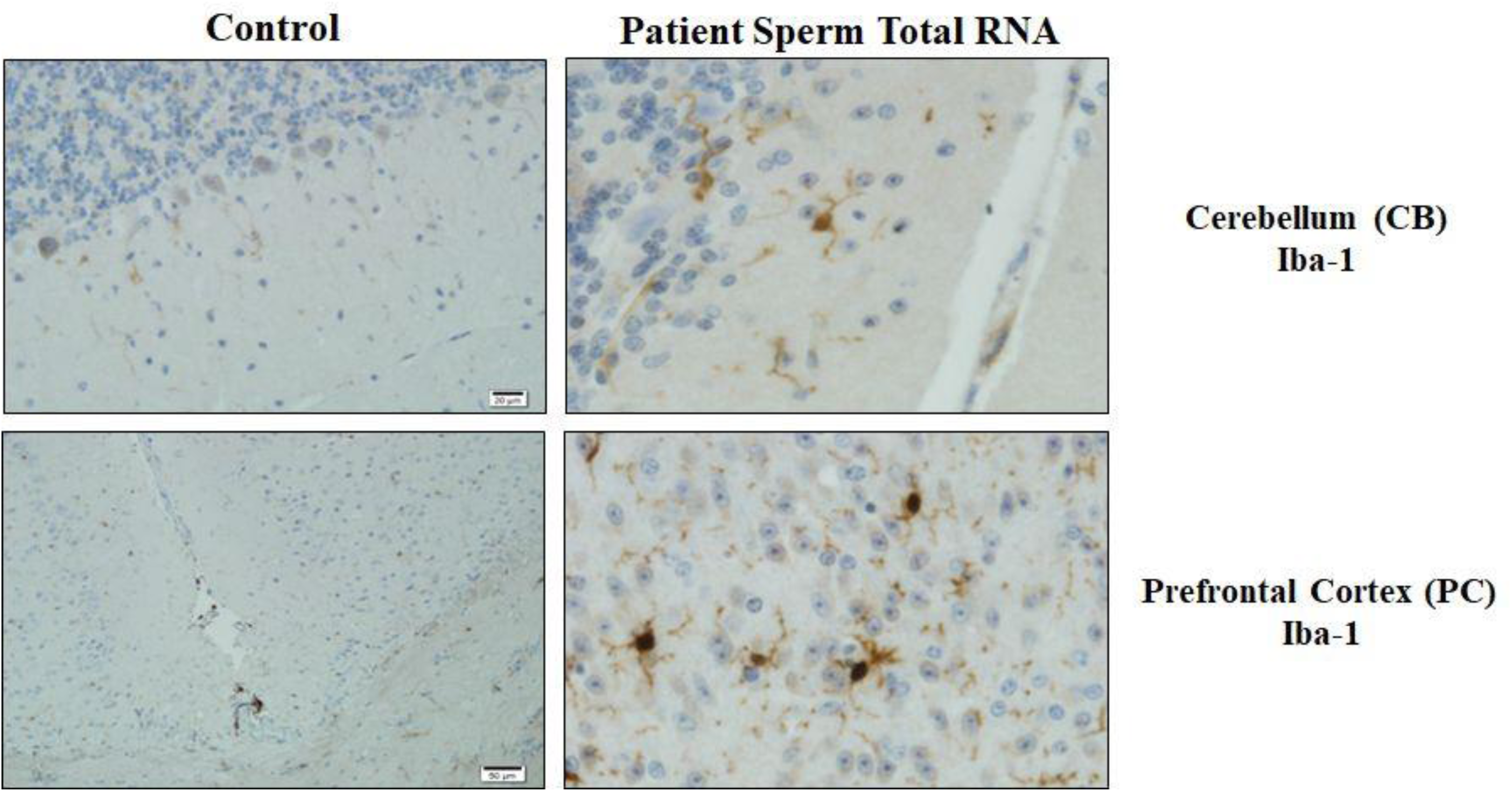
Visualization of expression levels in the singular cortex and with Iba-1 Ab staining and comparison of the patient father sperm RNA group with the control.

Although comparing the age of mice/or other animals to humans is not always appropriate, a young adult mouse could provide a first glimpse into the consequences of altered gene expression during embryonic development. The two-month period was necessary to obtain the comprehensive results from behavioural tests and histological analysis. Conducting, behavioural testing before puberty, especially with early separation from the mother’s littermate could potentially affect their behaviour.

In this study, we examined all sections of the brain with careful analysis of regions reported in previous studies. However, the most prominent observed alteration was related to glial cell and this effect was consistent between the VPA and exposed to RNA from autistic progenitor sperm or by microinjection of miR-499-3p. Other changes were also noted but were similarly observed in control exposed to human sperm RNA. The characterization of trans-species (human sperm) RNA-induced changes in mice can reveal specific RNA-mediated epigenetic signals. Our study not only confirmed the previously observed ASD-like behaviour in the 2-month-old mouse model, but also implicated RNA alterations in ASD development. Autism-linked miRNAs were down-regulated in the blood, hippocampus and sperm of two-month-old mouse models; along with corresponding changes in behaviour. This suggests a potential association between altered miRNAs expression markers in blood and changes in glial cells histology and phenotype. Indeed, as recently reported the glial cell signaling pathway was one of the most affected pathways. The alteration of glial cells accompanied by epi-molecular and phenotypic changes, align with early developmental stages and the critical period theory of ASD. The identification of glial cells in the mouse model is particularly important, as it supports that ASDs may be influenced by a mechanism involving the regulation of synaptic formation and function by glial cells.

Normalization of molecular functions at an early stage and finding an appropriate timing along with behavioural intervention, could be an effective therapy for ASD. The present study revealed that VPA mouse models exhibit glial, epi-molecular and behavioural phenotypes that are also induced by sperm RNAs from the father of a child with ASD. In particular, the comparable concordant RNA activity between mouse (VPA) and human ASD progenitor suggests that the mouse model is a suitable biological system for testing pharmacological treatments for human ASD.

Our previous studies on autistic behavior coupled with the findings morphological alterations in glial cells presented here, reaffirms that mice with ASD have continue to serve as a practical model for translational study. Furthermore, this study opens up several avenues for exploration. Firstly, routine testing of father’s sperm from individuals with different etiologies and symptoms could be considered. While the cellular, molecular and phenotypic changes in the mouse model of ASD in this study are similar with both human and mouse sperm RNAs in idiopathic ASD, it further validates the robustness of this model. Secondly, we highlight and confirm with the literature the roles of non-neuronal components such as astrocytes and microglia, whose accumulating evidence suggests involvement in the pathogenesis of ASD. Thirdly, the detection at ASD-related markers at two months of age in the mice found in this study allows for early screening diagnosis and treatment. That is particularly valuable as medications intended for newborns and children generally carry unwanted side effects and their effectiveness may not be fully established.

Nonetheless, this study indicated the crucial importance of RNA induced phenotypes in ASD. These results could contribute to future studies on how to track RNA-mediate diseases and plan new therapeutic approaches for ASD early in life^23^

## Supporting information

miRNAs' sequences for determination of miRNA expression levels

Morphological examination of the hippocampus in the group injected with miR-499a-5p reverse microinjection.

Histological examination of the hippocampus in different groups

Histological examination of different tissues in the group injected with 0.9% saline.

Histological examination of different tissues in male mice injected with 300 mg/kg VPA.

Histological examination of different tissues in the group injected with 400 mg/kg VPA.

Histological examination of different tissues in the group injected with 0.9% saline.

Demonstration of histological changes in female samples of the 500 mg/kg VPA group.

Examination of neuronal changes in cells using fluorescent staining in different groups and tissues.

Demonstration of the changes occurring as a result of histological and fluorescent staining in groups injected with 500 mg/kg VPA.

Representation of neural changes in hippocampus and striatum tissues in different animals in the 400 mg/kg VPA injected group with fluorescent stainin

Representation of neural changes in hippocampus and striatum tissues in different animals in the 300 mg/kg VPA injected group with fluorescent stainin

## Acknowledgments

This work was made possible by a grant to Y. Ozkul from Tübitak, 1010 (EVRENA project ID 112S570) and grant 2019-2020 “La Fondation Nestlé France” to Minoo Rassoulzadegan has participated to this work.

## Author contributions

All authors have approved the revised submitted version. M.R and Y.O. has organized and obtained financial support. Z.Y.Ş has designed and performed microRNAs analysis on the RNAs samples, behavioral analysis, data analysis and figure preperation. K.K.B has performed transferring human sperm RNA to mouse embryo and article preparation. E.M. has performed mouse behavior analysis, sample preparation for histological analysis. F.B, and Z.D. performed pathological and immunohistochemical analysis. S.T., M.R and Y.O has created project hypothesis, data interpretation, literature reviewing and manuscript preparation.

## Declaration of interests

Te authors declare no competing financial and/or non financial interests in relation to the work described.

## Supplementary Table and Figure Legends

**Supplementary Table 1:** Oligonucleotid’s sequences for microinjection

**Supplementary Table 2:** miRNAs’ sequences for determination of miRNA expression levels.

**Supplementary Figure 1.** Morphological examination of the hippocampus in the group injected with miR-499a-5p reverse microinjection.

**Supplementary Figure 2.** Histological examination of the hippocampus in different groups **A.** Control group, **B.** Patient father sperm RNA group, **C.** miR-499a-5p reverse group and **D.** F0 500 mg/kg VPA group.

**Supplementary Figure 3.** Histological examination of different tissues in the group injected with 0.9% saline. **A.** Morphological examination of striatum tissue, **B.** Morphological examination of hippocampus tissue.

**Supplementary Figure 4.** Histological examination of different tissues in male mice injected with 300 mg/kg VPA. **A-B.** Examination of morphological changes in different regions of the striatum tissue, **C.** Demonstration of morphological changes in the hippocampus.

**Supplementary Figure 5.** Histological examination of different tissues in the group injected with 400 mg/kg VPA. **A.** Morphological examination of striatum tissue, **B.** Morphological examination of hippocampus tissue.

**Supplementary Figure 6.** Histological examination of different tissues in the group injected with 0.9% saline. **A.** Morphological examination of striatum tissue, **B.** Morphological examination of hippocampus tissue and **C.** Morphological examination of cerebellum tissue.

**Supplementary Figure 7.** Demonstration of histological changes in female samples of the 500 mg/kg VPA group. **A.** Examination of morphological changes in the prefrontal cortex, **B.** Examination of morphological changes in the striatum cortex, **C.** Examination of morphological changes in the hippocampus, **D.** Examination of morphological changes in the cerebellum

**Supplementary Figure 8.** Examination of neuronal changes in cells using fluorescent staining in different groups and tissues. **A.** Demonstration of changes in the striatum in the saline group, **B.** Demonstration of changes in the hippocampus in the saline group, **C.** Demonstration of changes in the striatum in the 500 mg/kg VPA group, **D.** Demonstration of changes in the hippocampus in the 500 mg/kg VPA group.

**Supplementary Figure 9.** Demonstration of the changes occurring as a result of histological and fluorescent staining in groups injected with 500 mg/kg VPA. **A.** Demonstration of morphological changes in the hippocampus with histological staining. **B.** Demonstration of neural changes in the striatum with fluorescent staining.

**Supplementary Figure 10.** Representation of neural changes in hippocampus and striatum tissues in different animals in the 400 mg/kg VPA injected group with fluorescent staining.

**Supplementary Figure 11.** Representation of neural changes in hippocampus and striatum tissues in different animals in the 300 mg/kg VPA injected group with fluorescent staining. **A.** Demonstration of changes in the striatum, **B.** Demonstration of changes in the hippocampus.

## METHODS

### Ethics Statement

This study was approved by the Hospital Ethics Committee and authorizations from all the patients and the participating relatives were obtained by signing an informed consent form. All parents gave written informed consent before participation (09-20-2011 committee number: 2011/10). All research was performed in accordance with the relevant guidelines and regulations (Erciyes University animal ethics committee 04-11-2012 12/54).

## 1. Patient Selection Criteria

This study was approved by the Hospital Ethics Committee of Erciyes University School of Medicine. A detailed description of the study was given to all participants and their parents before their enrollment. All parents gave written informed consent before participation. The diagnosis was made by a multidisciplinary team (comprising an experienced child psychiatrist, a pediatric neurologist, and a genetic specialist), according to the criteria of the Diagnostic and Statistical Manual, Fourth and Fifth Edition, Text Revision (DSM-IV-TR; American Psychiatric Association, 2000 and DSM-V; American Psychiatric Association, 2013) criteria, using Childhood Autism Rating Scale (CARS). From the Turkish cohort of 37 families including one or more children with behavior disorders (45 subjects altogether)^9^, we were interested in a family with two affected children. A father from two separate marriages has multiple children and one of each bed develops different psychological disorder. We have previously shown that all of them (six miRNAs) are therefore already altered in their father’s sperm. Because of this reason, we planned to microinjected father’s sperm RNA to mouse embryos. For control, heathy control father’s sperm RNA were taken.

## 2. Human sperm RNAs preparation

The same microRNAs reduce in blood of autistic patients were also reduced in the sperm RNAs of the autistic children father^9^. To follow any effect from the human sperm RNAs in mice, we have prepared total sperm RNAs from two men one control and second from the father of the autistic children. Totals Sperm RNAs were prepared by standard Trizol extraction techniques (see methods). RNAs for microinjection into fertilized mouse eggs were adjusted to 0.2ng/microliter from control sperm and father of children with autism (see table 1 for the number of animals and concentration of RNAs and microRNAs). Mice born from microinjection were designed SH*(Sperm Human SHA* for Autism) and miR*. In the group microinjected with human sperm RNA, there is no difference between the groups in terms of behavior or any miRNAs expression (see Supplementary Figure xx). There is some effect of human sperm RNAs in the mouse in behavior tests but it seems to be specific to human sperm RNAs in general rather than a specific signal from the sperm RNAs of the father of children compared to control human sperm RNAs.

## 3. Mouse husbandry

Mouse were maintained according to the European regulations for the care and use of research animals. The genetic backgrounds are B6D2 F1 and Balb/c.

## 4. RNA microinjection

Oligoribonucleotides synthetic miRNA were adjusted to a concentration of 1 µg/ml and microinjected into normal Balb/c fertilized eggs according to established transgenesis methods^12^ Oligoribonucleotides were obtained from Eurofine (sequences provided in Supplemental Table S1).

Briefly, for collecting embryos, female and male mice were mated at 3 pm. The day after, mated females were checked for the vaginal plug at 8 am and vaginal plug (+) females mice were selected. Females were sacrificed that afternoon, and their oviducts were enclosed in M2 medium (Sigma, Germany) and embryos were collected under a bino-microscope (Leica, Germany) and were transferred into M16 medium (Sigma, Germany) with a mouth pipette and incubated at 37°C in 0.5% CO2 (Panasonic, Japan). To capture the pronuclei at the most prominent stage, at 2-6pm we microinjected oligonucleotides (1-5 ng/µl, see in Supplemental Materials Table S1) with a glass pipette into the male pronucleus of embryos under an inverted microscope (Nikon, Japan). After the microinjection, the embryos were incubated at 37°C in 0.5% CO2 (Panasonic, Japan). Embryos that died after microinjection under a stereo microscope (Leica, Germany) were separated and surviving embryos were transferred into M2 medium (Sigma, Germany). Living embryos were transferred to the foster mother (healthy female with a vaginal plug after mating with vasectomized male mice), under anesthesia with a mouth pipette into the oviduct. Three weeks later F0 generation pups, which were born 21 days after the procedure, were separated from their mothers according to their genders and transferred to new cages and subjected to behavioral and molecular tests with other groups at two months of age.

## 5. Behavior tests

Behavioral experiments were started when miRNA microinjected and control group mice were 2 months of age. Each mouse underwent a single test daily between 10:00 and 16:00. Only males were tested sequentially on the same day in separate sub-sessions to allow room ventilation and cleanup. The testing tools were cleaned between trials with 70% ethanol and aerated before use. Experiments were videotaped and analyzed offline. Sociability, social-preference and object-recognition and tail suspension tests were analyzed using “EthoVision 9” software (Noldus, Wageningen, Netherlands). Marble burying tests were analyzed manually by an observer blind to the group of the mice.

### 5.1. Novel Object Recognition Test

A mouse-size object and a second identical object were placed in a square box with an open top with lines dividing it at the base and a wall enclosing it. A mouse-sized object and a second identical object are placed. The mouse’s proximity to the objects and the number of visits were counted on the first day. On the second day, one of the objects was replaced with a new object. The discriminating index, the number of times the mouse approached both unfamiliar and recognized objects, and their proximity were all measured. The data obtained is defined as the discrimination index when the difference between the total time spent with the new object and the total time spent with the familiar object is divided by the total duration and multiplied by 100. The proportion of the mice’s overall interaction time with the new object is measured. The learning and memory abilities of mice are examined in this experiment. Because mice have an innate desire for novelty, it was anticipated that they would spend more time with the novel object to discover more about it ^25^

### 5.2 Social interaction Test

The Social Test assesses cognition in mouse models of Central Nervous System (CNS) disorders by evaluating general sociability and interest in social novelty. Since they are sociable creatures, rodents normally prefer to spend more time with other rodents (sociality) and are more likely to approach a stranger than a friend (social novelty). The rectangular, three-compartment box used in Crawley’s friendliness and social novelty test con-sists of a rectangular three-compartment box. Every room is 19 × 45 cm and is a system with partitioned walls and a central chamber made of clear plexiglass that allows open access to every area^26^. A mouse is initially placed into a middle compartment for 5 minutes while the other compartment was left empty. Under the parameters found to be connected to the mouse in the chamber, data were generated using the EthoVision system, and statistical analysis was performed along with comparisons to the control group.

### 5.3. Marble Burying Test

The Marble Burying test is commonly used to evaluate rodent neophobia, which includes shyness around unfamiliar objects, anxiety and obsessive-compulsive or repetitive behaviors^27^. We positioned the bedding that we usually use to care for our mice empty cage at the height of 5 cm in an empty cage. There are five rows of four marbles each, totaling twenty marbles. The experimental mouse was let out of one corner of the cage and given 30 minutes to roam around. The mouse was removed from the cage after 30 minutes, and the number of balls discovered under the bedding was tallied and recorded. The number of embedded balls was used for statistical analysis.

### 5.4. Tail Suspension Test

The mouse suspended by the tail test is a model experiment model of autism on ob-serving that after initial escape-directed movements, mice develop a sedentary stance when placed in an unavoidably stressful situation. The stressful situation during tail-hanging includes the hemodynamic stress of hanging, and autistic models seem to have reduced mobility and escape abilities^28^. The experimental setup was designed to simultaneously test three different mice. Thick cardboard-like sheets measuring 25 cm high were cut and placed between the mice so that they could not see each other. Since the mice are white in color, a black background background was used. Twelve cm-long tapes were cut and hung in the experimental setup by sticking them to the tail ends of mice in such a way that their tails would not be damaged. By watching recordings with a video camera, the mobility and immobility time of mice were calculated for six minutes. The immobility time was used for statistical analysis.

After the behavioral experiments of the F0 generation were completed, two females and one male were mated to obtain the F1 generation. When obtaining the F1 generation, the F0 generation and the control group were sacrificed and blood, hippocampus and sperm samples were taken. For the F1 generation, behavioral experiments were conducted when they were 2 months old. The F1 generation was only studied for behavioral changes and molecular analyzes were not performed.

## 6. RNA isolation From Tissue

Sperm and blood samples were taken into 500 µl Purezol (Biorad, USA, CA, Cat No: 7326890) Subsequently, total RNA isolation was carried out according to the manufacturer’s instructions.

## 7. cDNA Preparation and Quantitative Real-Time Polymerase Chain Reaction (qRT-PCR)

Isolated RNA samples were reverse-transcribed into cDNA in 20 μl final reaction volumes using miScript II RT Kit (Qiagen,,Germany) as specified in the manufacturer’s protocol. Reverse transcription was performed using the SensoQuest GmbH Thermal Cy-cler (Göttingen, Germany). cDNA samples were kept at −80°C until PCR analysis.

QRT-PCR was performed by using miScript SYBR® Green PCR Kit (Qiagen, Hilden, Germany Cat No: 218073) with the high-throughput Light Cycler 480 II Real-Time PCR system (Roche, Germany, Mannheim). cDNA samples were diluted with Nuclease Free Water (1:5). The reaction was performed according to manufacturer’s instructions. About 10 μl Syber Green Master Mix, 2 µl 10x Universal Primer, 2 µl primer assays and 4 µl nuclease free water mixed and pipetted into a 96 well plate as 18 µl and 2 μl of 1:5 diluted cDNA was pipetted into each well and mixed. The Real-Time PCR step was performed by using the Light Cycler 480 II Real-Time PCR system with the following protocol: thermal mix followed by the Activation step at 95°C for 15 min, then a denaturation step at 94°C for 15 sec followed by an annealing step at 55°C for 30 sec followed by an extension step at 70°C for 30 sec. After activation step, all steps were carried out for 40 cycles. Data normalization was performed using the 2^-ΔΔCT^ method with U6 internal control

## 8. Data Analysis

After the results were obtained, experimental groups and control groups comparisons were made. The compliance of the data to normal distribution was evaluated by the histogram, q-q graphs and Shapiro-Wilk test. Statistical analysis was performed by two-tailed, one-way analysis of variance (ANOVA), uncorrected Fisher’s least significant difference (LSD). Kruskal Walls, student t-test and Mann Whitney-U test depending on whether the data showed normal distribution or not. Data were analyzed using SPSS version 22 (IBM, USA) and Graph-Pad Prism 8.0 software. Results with p values <0.05 were considered statistically significant. Data are expressed as the mean with SD.

## 9. Immunochemistry

Different mice were deeply anesthetized and transcardially perfused with 4% paraformaldehyde. Series of, x-mm sections were incubated in a solution of mouse anti- (1:1000,), rabbit anti- (1:500,), goat anti- (1:500,) and rabbit anti-GFAP (1:500,) for 48h at 4C. After several rinses in phosphate-buffered saline of: x-conjugated highly cross adsorbed donkey anti-goat, (all at 1:500; Life Technologies) in PBST. Sections were rinsed intensively and mounted onto slides, coverslipped using Mowiolmixed with Hoechst (1:10 000; Life Technologies) and stored at 4°C.

## Notes

### Competing Interest Statement

The authors have declared no competing interest.

