## Supplementary figures and images for "RNA activities of trans species: RNA from the sperm of the father of an autistic child programs glial cells and behavioral disorders in mice"

### Demonstration of histological changes in female samples of the 500 mg/kg VPA group.

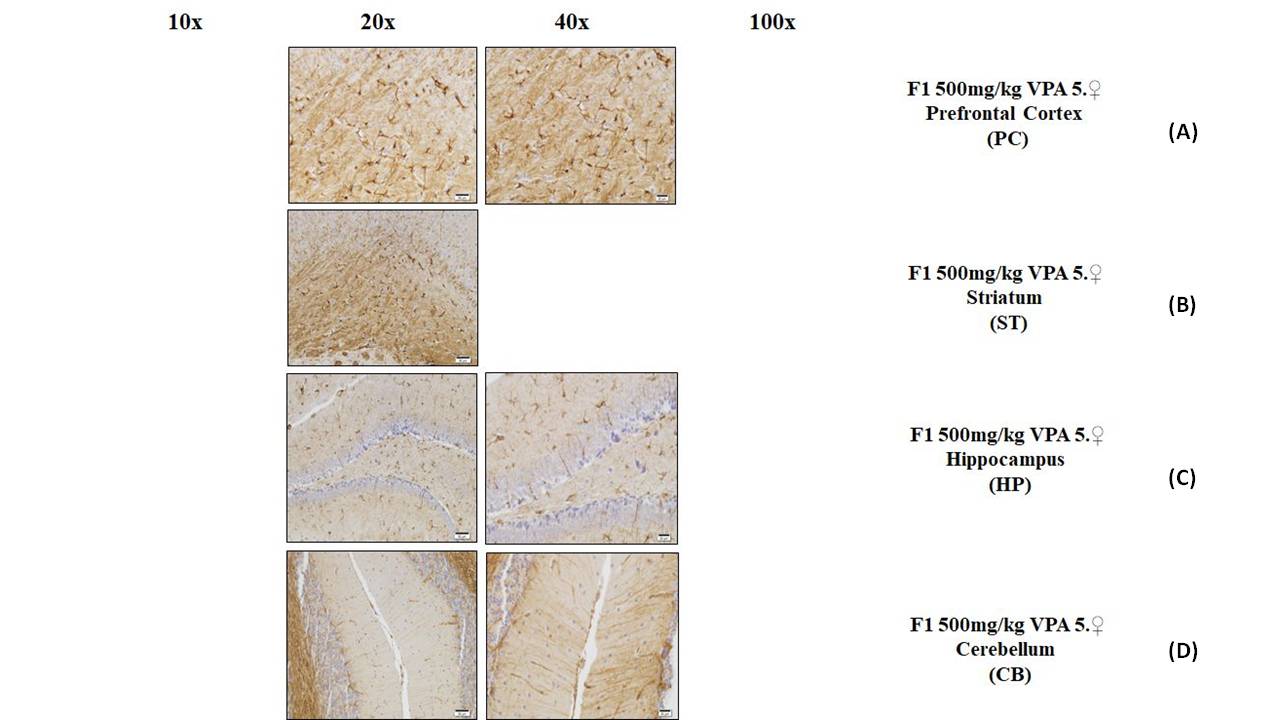

### Demonstration of the changes occurring as a result of histological and fluorescent staining in groups injected with 500 mg/kg VPA.

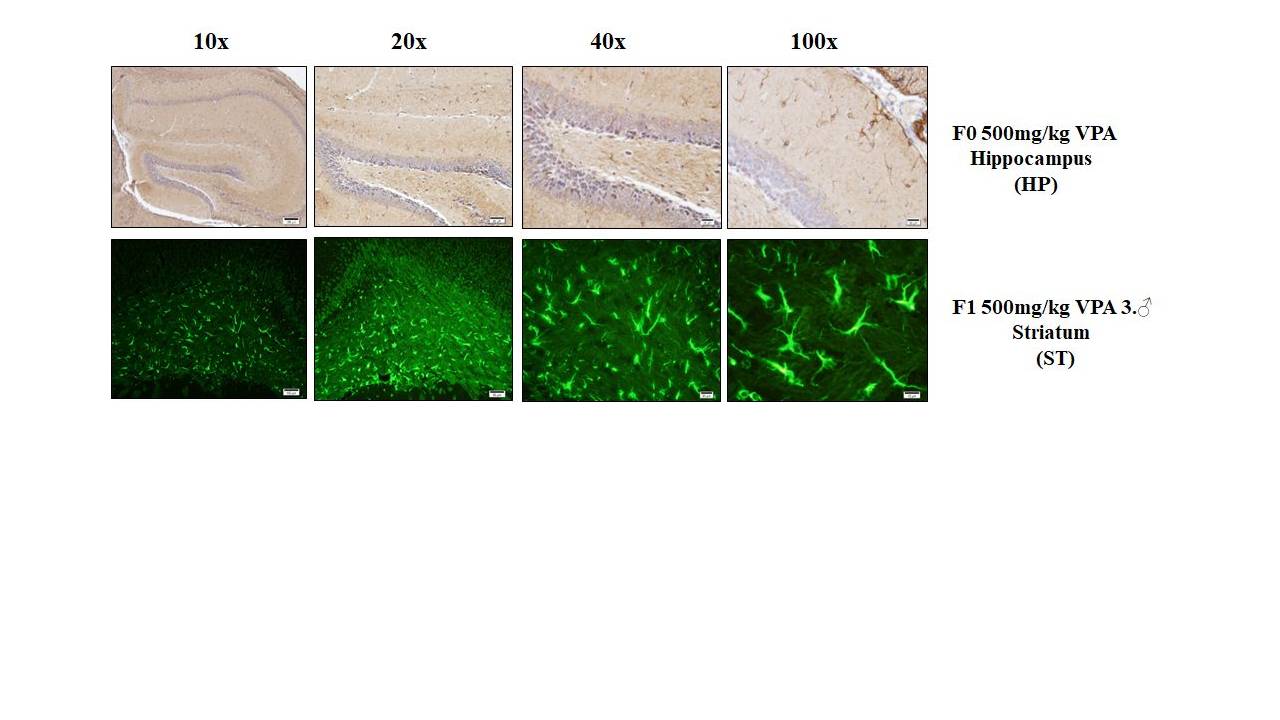

### Examination of neuronal changes in cells using fluorescent staining in different groups and tissues.

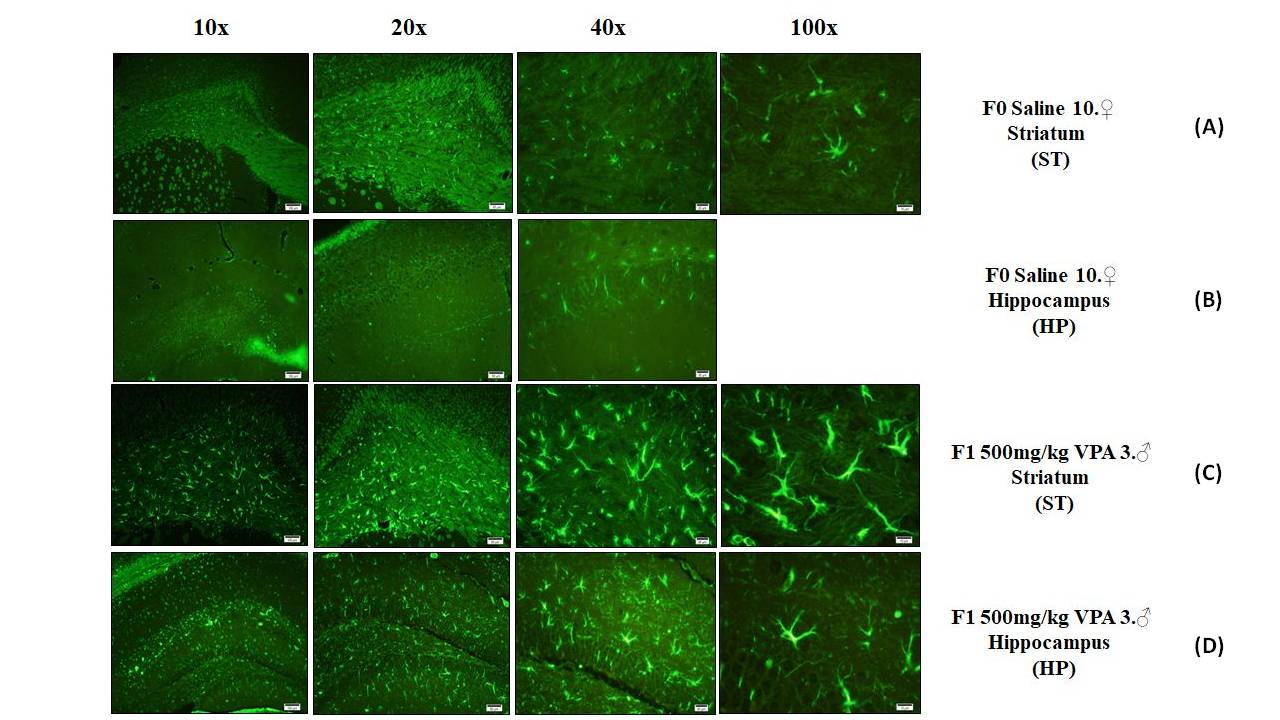

### Histological examination of different tissues in male mice injected with 300 mg/kg VPA.

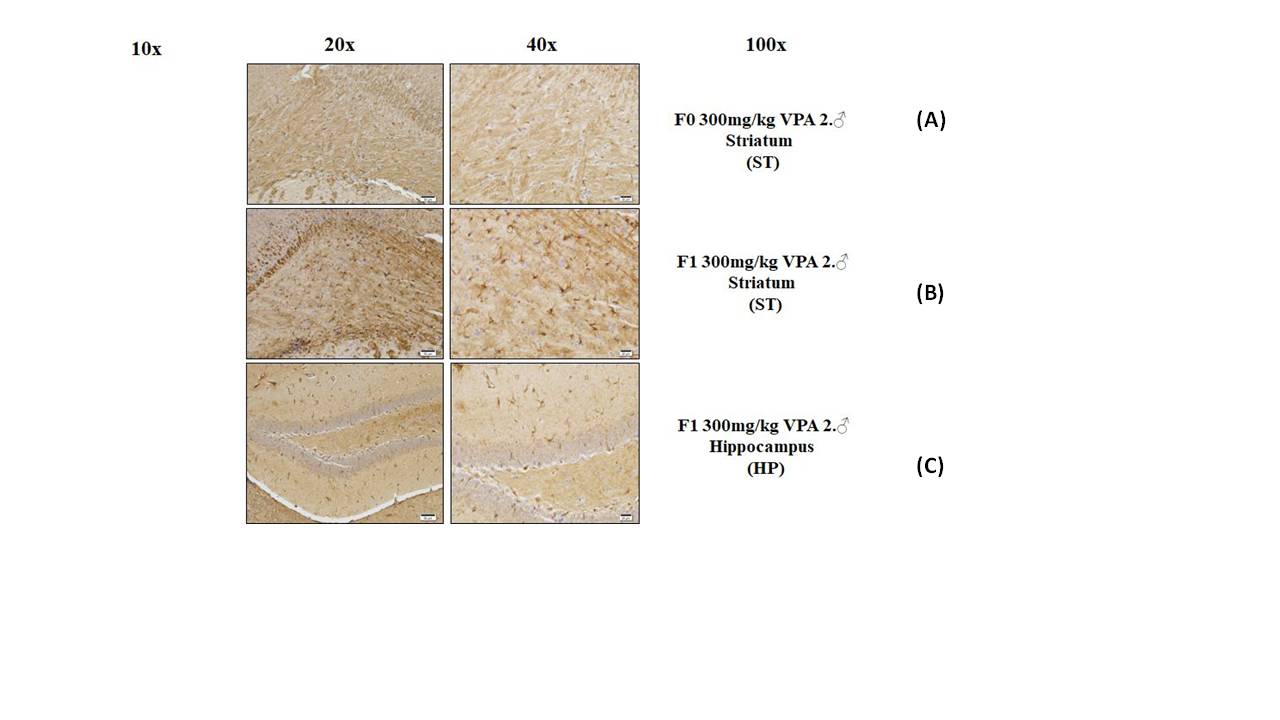

### Histological examination of different tissues in the group injected with 0.9% saline.

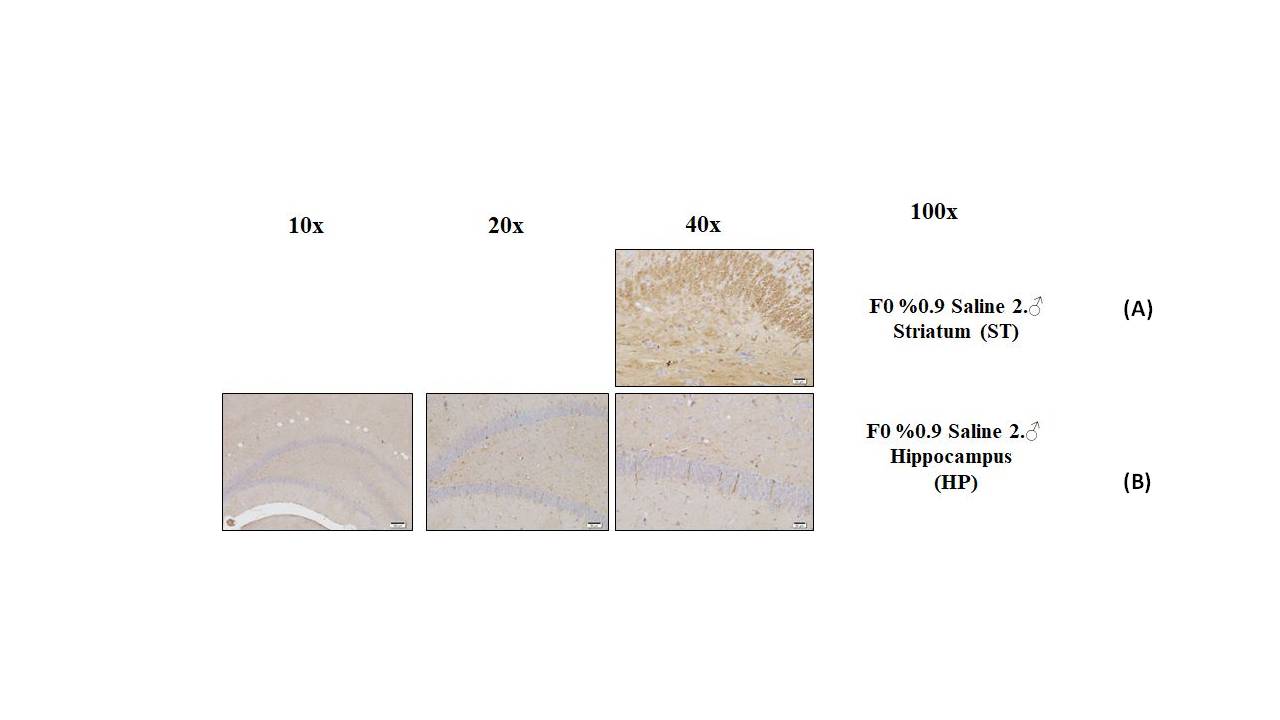

### Histological examination of different tissues in the group injected with 0.9% saline.

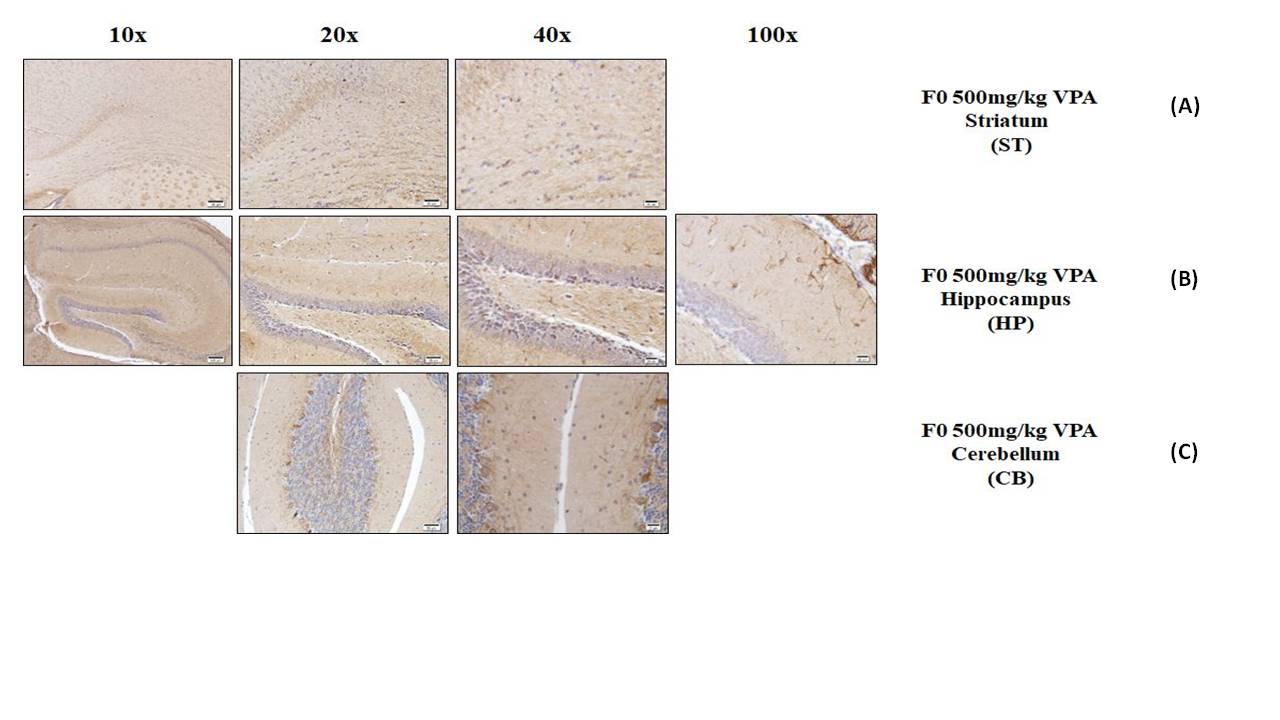

### Histological examination of different tissues in the group injected with 400 mg/kg VPA.

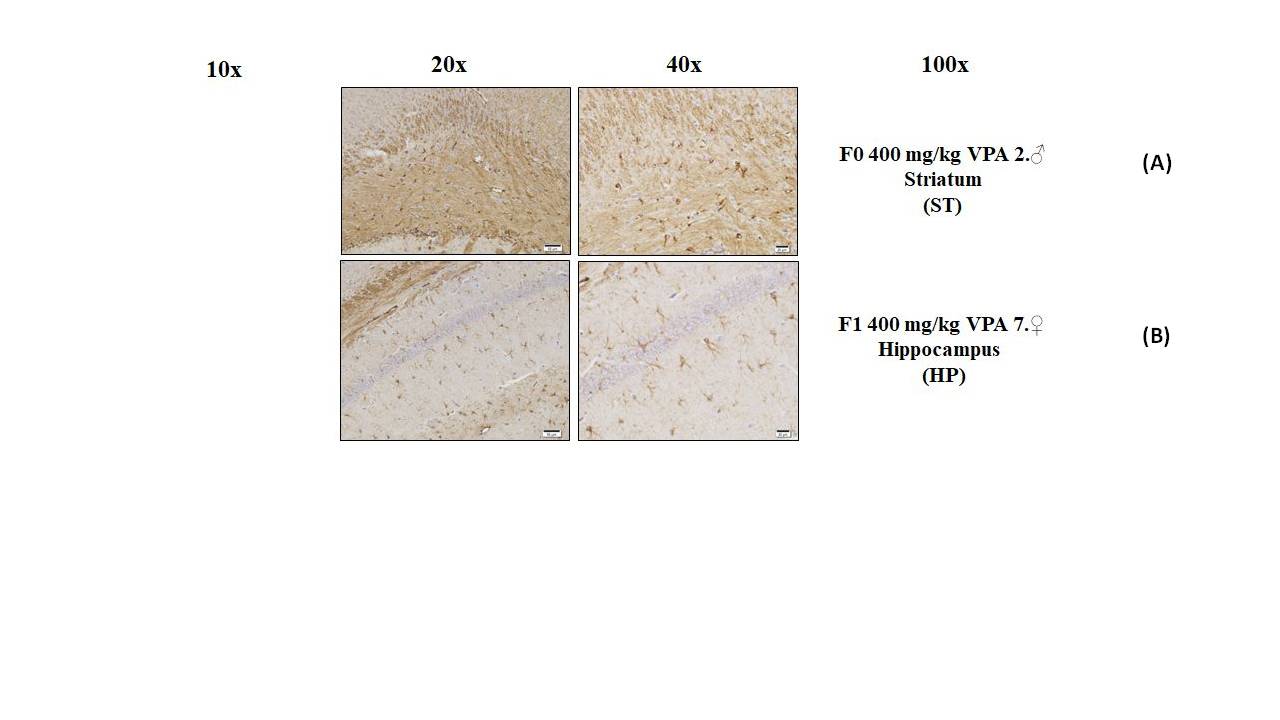

### Histological examination of the hippocampus in different groups

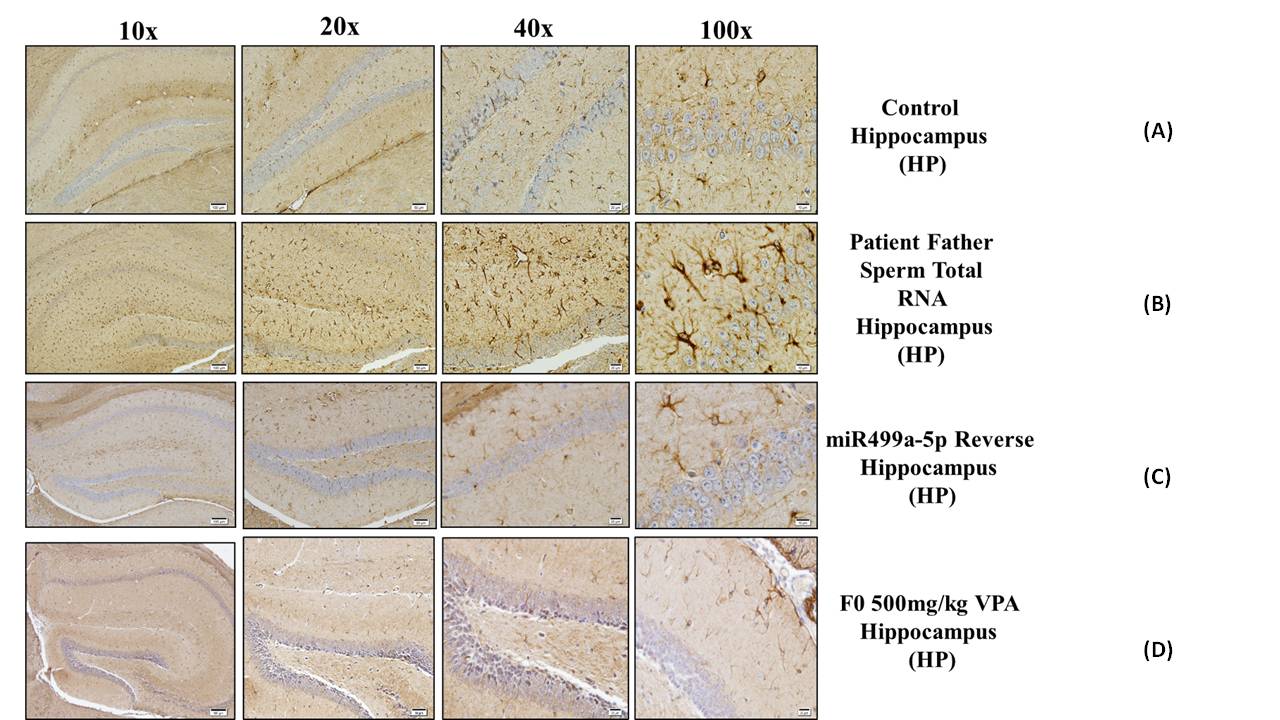

### miRNAs' sequences for determination of miRNA expression levels

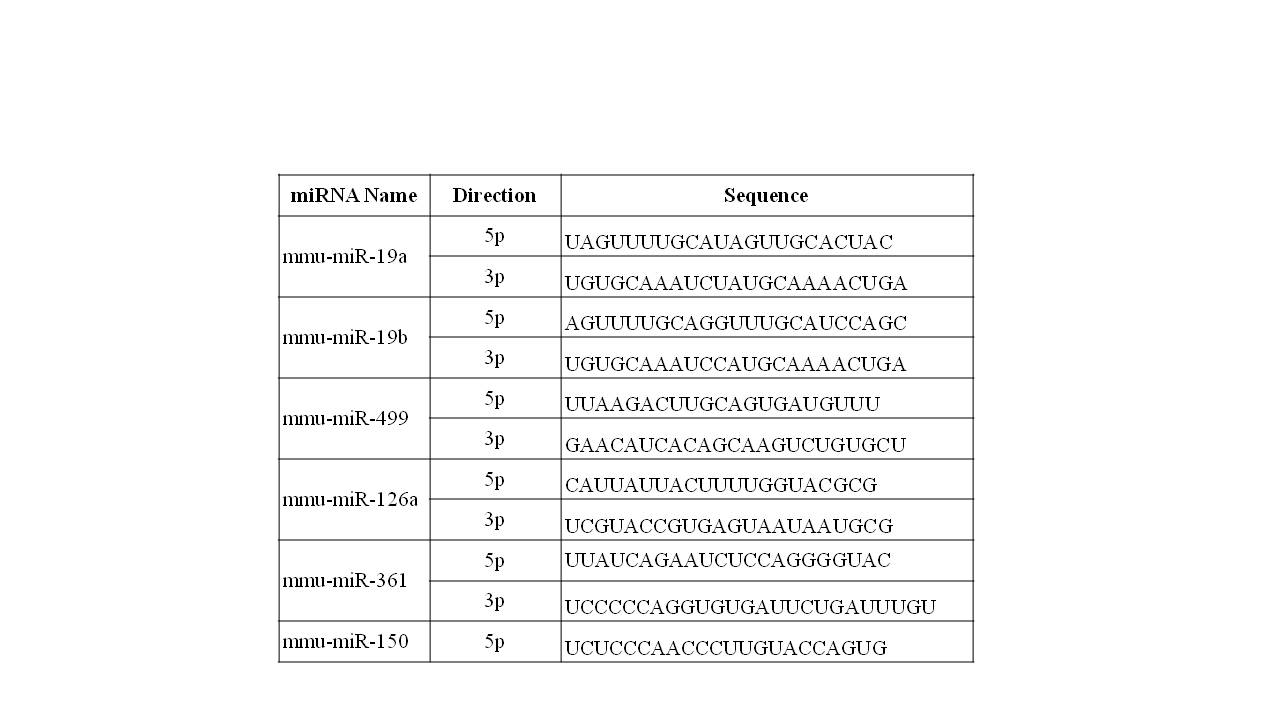

### Morphological examination of the hippocampus in the group injected with miR-499a-5p reverse microinjection.

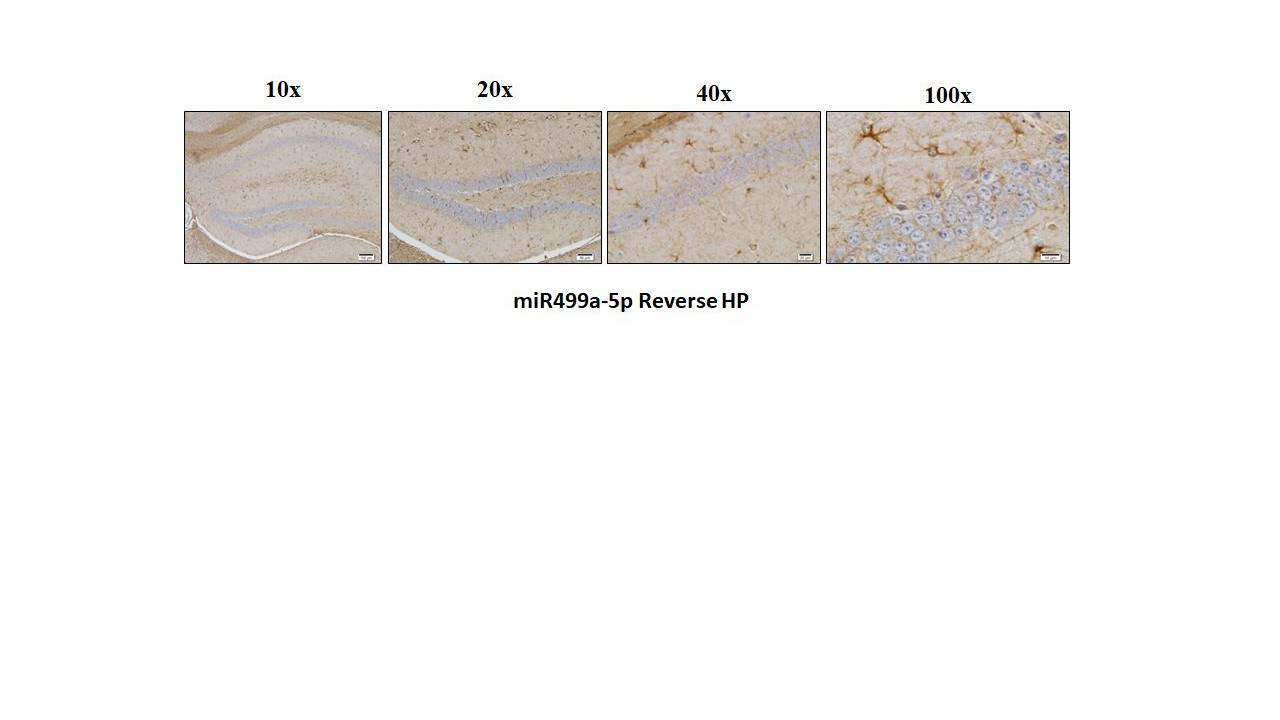

### Representation of neural changes in hippocampus and striatum tissues in different animals in the 300 mg/kg VPA injected group with fluorescent stainin

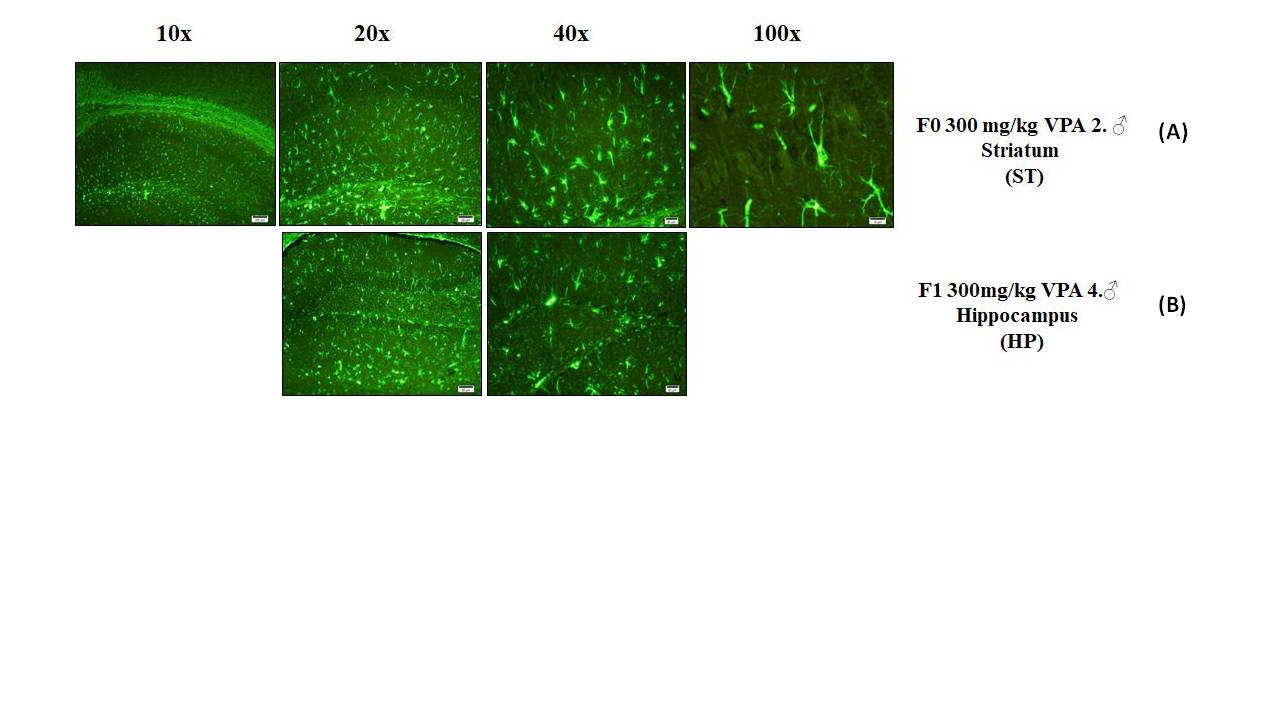

### Representation of neural changes in hippocampus and striatum tissues in different animals in the 400 mg/kg VPA injected group with fluorescent stainin

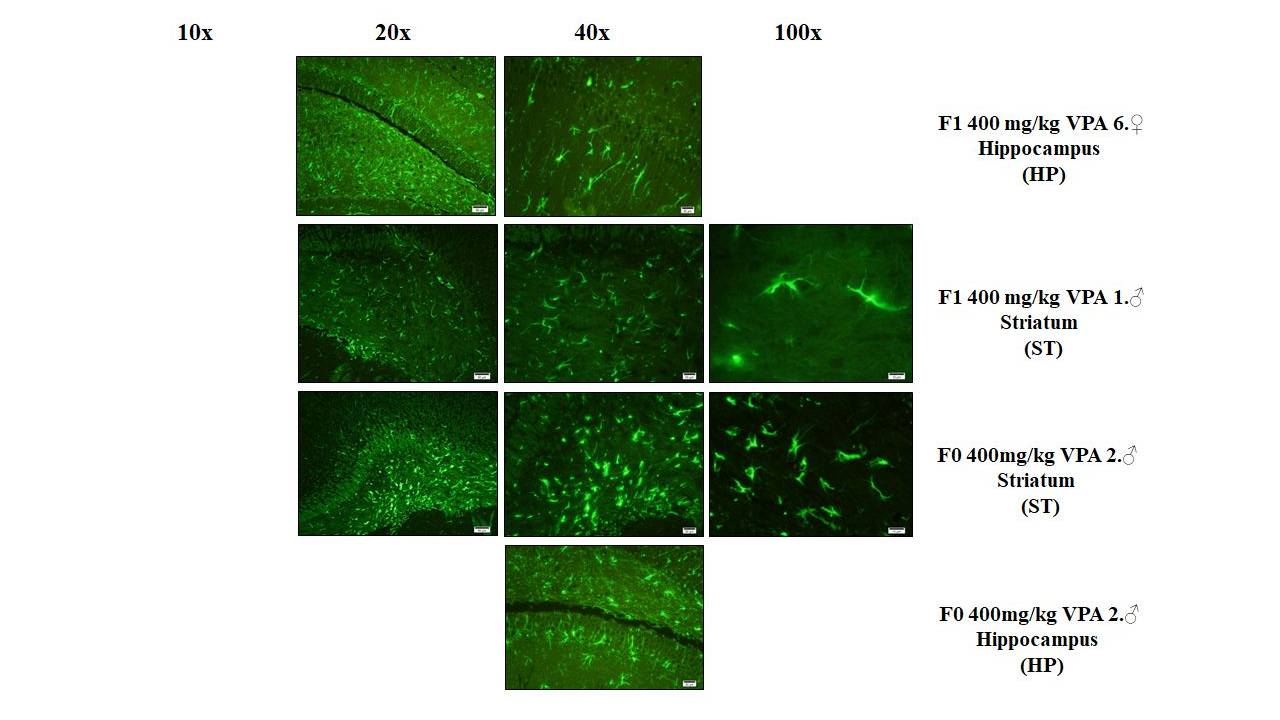

### Supplemental Data 1

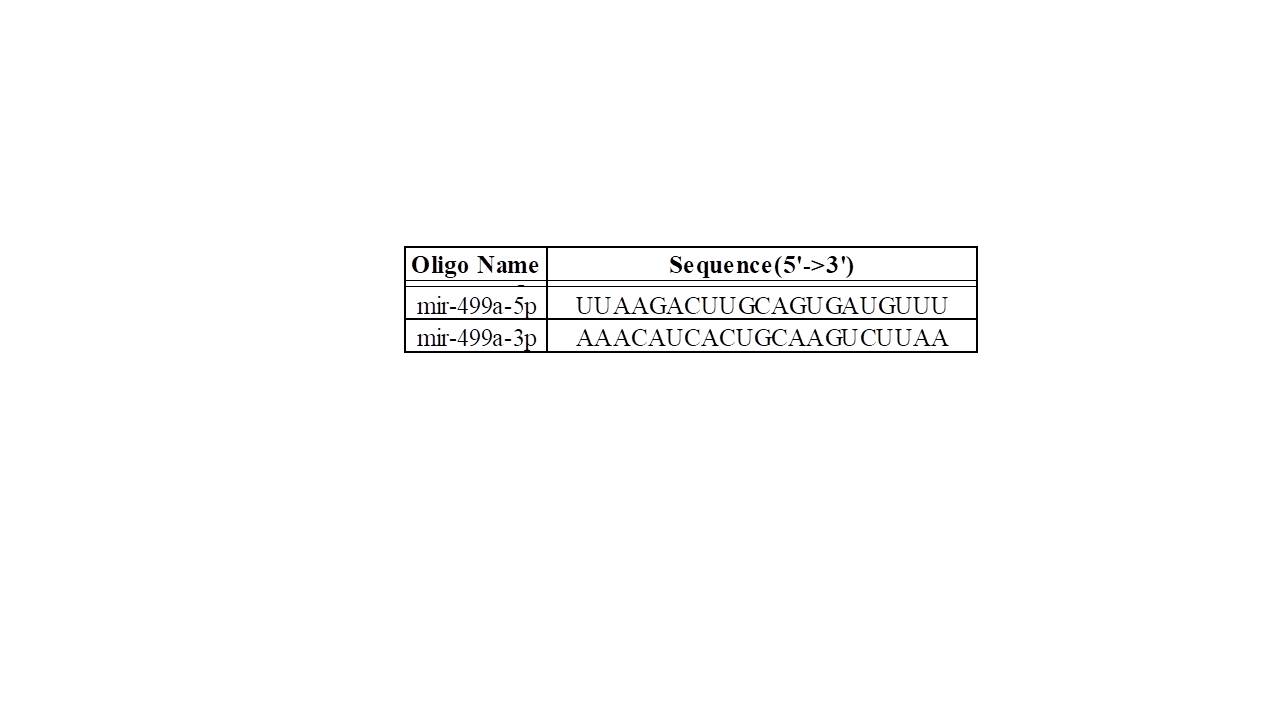
